# Night-time control of carbohydrate availability in grasses differs radically from that in Arabidopsis

**DOI:** 10.64898/2026.06.02.729567

**Authors:** Matthias Thalmann, Alison M. Smith

**Affiliations:** John Innes Centre, Norwich NR4 7UH, United Kingdom

## Abstract

Grass leaves reportedly accumulate sucrose during the day then export it to other organs at night. The control of export at night is not understood. By contrast, Arabidopsis leaves accumulate starch then convert it to sucrose for export at night. Control of starch mobilisation ensures a constant sucrose supply and exhaustion of starch around anticipated dawn. We found that that 41 species of pooid grasses differed widely in the ratio of sucrose to starch accumulation and night-time depletion (turnover) in leaf blades, regardless of growth conditions. Sucrose turnover exceeded starch turnover in most species, but in a minority of species the two were similar. In eight species spanning this diversity, the rate of sucrose depletion fell through the night. In six of the species, all with higher sucrose than starch turnover, starch depletion at night was initially slow. Later, it increased to a rate that exhausted starch around dawn. The initial lag was absent and starch depletion was linear throughout the night in the other two species, both of which had similar sucrose and starch turnover. We suggest that underlying control of starch mobilisation in grass leaf blades resembles that in Arabidopsis leaves. The initial lag in starch depletion in high-sucrose species may result from suppression of starch degradation by sucrose, acting via the sucrose signalling metabolite Tre6P. The large diel fluctuations in sucrose export from grass leaf blades may be dampened by exchange with dynamic pools of sucrose in leaf bases and sheaths prior to export to other organs.

## INTRODUCTION

The aim of this work was to shed light on diel patterns of carbohydrate storage and depletion in leaves of grasses in the Pooideae, a C_3_ subfamily of the Poaceae that contains the cereals wheat, barley, oats and rye and many temperate pasture and amenity grasses.

Growth of Arabidopsis plants during the night is sustained by consumption of leaf starch reserves accumulated by photosynthesis during the preceding day. In carbon-limited conditions (typically 12 h or less of light per day) the rate of starch consumption is constant through the night, and is such that reserves are exhausted almost exactly at dawn. This pattern of consumption maximises the fraction of photosynthate available for diel growth and persists under a range of environmental conditions (Graf et al., 2010; Stitt and Zeeman, 2012; Scialdone et al., 2013; Sulpice et al., 2014; Pilkington et al., 2015; Annunziata et al., 2017; Mengin et al., 2017). Mutations that reduce the rate of starch consumption at night reduce growth rate in carbon-limited conditions (e.g. Fulton et al., 2008; Feike et al., 2016; Chew et al., 2022). The mechanism that controls starch consumption in Arabidopsis leaves requires both the circadian clock and the accumulation of reserves during the day in the form of starch granules (Scialdone et al., 2013; Pyl et al., 2012, Ishihara et al., 2022). However leaves of many species of plants, including members of the Pooideae, accumulate reserves during the day predominantly in the form of sucrose rather than starch. Relatively little is known about how, and indeed whether, the rate of sucrose depletion in these species is controlled to maximise diel growth in carbon-limited conditions in a manner analogous to starch consumption in leaves of starch-accumulating species. (“Sucrose depletion” is used here to mean its loss at night, due to both local metabolism and export.)

Diel carbohydrate turnover in leaves of species in the Pooideae has been studied extensively in leaves of the closely-related cereal crops wheat (*Triticum aestivum*) and barley (*Hordeum vulgare*). Studies differ widely in the reported absolute and relative rates of diel carbohydrate turnover, probably reflecting the wide range of different growth conditions, sampling strategies, leaf and plant ages and genotypes employed. Nonetheless it is clear that both species accumulate large amounts of sucrose and smaller amounts of starch in their leaves during the day, and degrade both carbohydrates during the subsequent night (e.g. Gordon et al., 1980a, b; Sicher et al., 1984; Schnyder, 1993; Balaguer et al., 1995; Nie et al., 1995; Shaihk et al., 2000; Trevanion, 2000; Müller et al., 2018). Studies on other species in the Pooideae provide a generally similar picture. For example, diel sucrose turnover is reported to exceed starch turnover in leaves of *Phleum pratense* (timothy: Bertrand et al., 2008), species of *Poa* (Borland and Farrar, 1985), *Phalaris aquatica* (Ciavarella et al., 2000), and *Lolium perenne* (ryegrass: Cairns et al., 2002). Numerous other species in the Pooideae are also reported to have high levels of sucrose and generally lower levels of starch in their leaves (e.g. Sakai and Hayashi, 1973; Schnyder, 1993; Barbehenn et al., 2004), so it is reasonable to assume that that higher sucrose than starch turnover is a common feature of the subfamily. The model grass species *Brachypodium distachyon* appears to be an exception to this general picture: it is reported to have a greater diel turnover of starch than of sucrose in its leaves (Jensen and Wilkerson, 2017; Müller et al., 2018).

Detailed studies of carbohydrate turnover in barley leaves revealed that accumulation of sucrose during the day was high at high light intensity, and loss of sucrose in the subsequent night was exponential rather than linear, regardless of sucrose content (Gordon et al., 1980a; Müller et al., 2018). Gordon et al. (1980a) reported that loss of starch during the night was also non-linear. An initial lag in the onset of starch degradation was dependent on the light intensity at which plants were grown. End-of-day starch content was low and the lag was minimal in plants grown at low light intensity, but content was higher and the lag more prolonged in plants grown at high light intensity. Non-linear patterns of depletion of sucrose and starch at night have also been reported in wheat leaves (Trevanion, 2000).

Understanding diel carbohydrate turnover in leaves of the Pooideae is complicated by the accumulation of fructans in leaves of some members of this subfamily (Smouter and Simpson, 1988). Species in the core Pooideae (constituting 80% of species in the Pooideae including tribes Poeae and Triticeae; Zhang et al., 2022) contain genes encoding known fructosyltransferases with a characteristic conserved motif (Sandve and Fjellheim, 2010; Schubert et al., 2019). Leaves of many of these species contain fructans, although reported amounts are highly variable. Analyses of circumstances under which fructans accumulate indicate that expression of fructosyltransferase genes is induced when sucrose accumulates, for example in response to cold, drought, high light, long days, and low sink demand (e.g. Cairns et al., 2002; Thomas and James, 1999; Versluys et al., 2018; del Pozo et al., 2019). Conversion of sucrose to fructans may serve to prevent sucrose concentrations rising to levels that inhibit photosynthesis (Pollock, 1986). Species in tribes outside the core Pooideae, for example in the Brachypodiaceae and Stipeae, lack genes encoding the classic fructosyltransferases of the core Pooideae (Li et al., 2012).

There are contradictory reports about the extent of diel turnover of fructans in the core Pooideae. Diel fluctuation in fructan levels was reported for the flag leaf of wheat (Nie et al., 1995) and for leaves of young barley plants (Müller et al., 2018), but other studies found minimal diel fluctuation in fructan levels in leaves of barley (Gordon et al., 1982; Sicher et al., 1984; Farrar and Farrar, 1985; Koreleva et al., 1997, 1998; Barros et al., 2020), tall fescue (*Festuca arundinacea*), Italian ryegrass (*Lolium multiflora*), *Poa* spp. and *Phalaris aquatica* (Lechtenberg et al., 1972; Borland and Farrar, 1985; Ciavarella et al., 2000; de Souza et al., 2005). There are huge differences between studies in amounts of fructans reported for a given species, likely due to differences in growth conditions (including photoperiod, light intensity and temperature), plant and leaf age, sampling strategies, and the technical challenges of making quantitative measurements of oligofructans (Matros et al., 2019).

To obtain a general picture of diel carbohydrate turnover in the Pooideae we examined starch and sucrose levels at the start and end of the light period under carbon-limited conditions (10-h light periods) in 41 species giving a broad coverage of the subfamily, then chose a representative subset of species with different sucrose to starch ratios for further examination. We explored the diel patterns of starch and sucrose turnover and the impact of daylength and light level on carbohydrate levels. We also checked for diel fructan turnover under our growth conditions in species known to be capable of fructan synthesis. All measurements were made on material from the mid-point of the blade of the youngest fully expanded leaf, avoiding the basal zone in which carbohydrate turnover differs from that in the distal parts of the blade (Schnyder et al., 1988; Allard and Nelson, 1991; Shaihk et al., 2000; Xiong et al., 2015; Czedik-Eysenberg et al., 2016). We found greater sucrose than starch accumulation in the light in almost all species examined. In all species both sucrose and starch were both substantially depleted during the night. However, there were large differences between species in the ratio of sucrose to starch accumulated in the light, in the patterns of night-time depletion of these two products, and in the impact on carbohydrate turnover of light intensity and day length. At least part of this interspecific variation is probably of recent evolutionary origin.

## RESULTS

### Both sucrose and starch undergo diel turnover in species in the Pooideae

An initial survey of starch and sucrose levels at the end of the day (EoD) and the end of the night (EoN) used 41 species, representing six out of the 16 tribes in the subfamily Pooideae and including the four tribes with the largest numbers of species (tribes indicated on Figures 1 and 2; Zhang et al., 2022). Within the tribe Pooeae, which contains around 65% of the genera and 60% of the species in the Pooideae, our species selection represented 13 out of the 24 subtribes (shown on Figure 2), and the major subdivisions proposed for this tribe (Chloroplast Groups 1 and 2 defined by Soreng et al., 2017; Nuclear Groups 1 and 2 defined by Zhang et al., 2022). Plants were grown in a glasshouse under ambient conditions of light and temperature, with a 10-h light period achieved by automated movement of shelves into and out of a dark shed. Measurements of starch and sucrose were made on 1-cm sections taken from midway along the blade of the youngest fully expanded leaf of young, non-flowering plants, grown either from seed or from divisions of non-flowering stock plants. At each harvest, five or six leaf samples were taken per species at EoD and EoN, each from a different plant. Five batches of between four and 11 species, selected without phylogenetic bias, were grown and sampled between April and September (sampling dates indicated by coloured circles on Figures 1 and 2).

**Figure 1.**
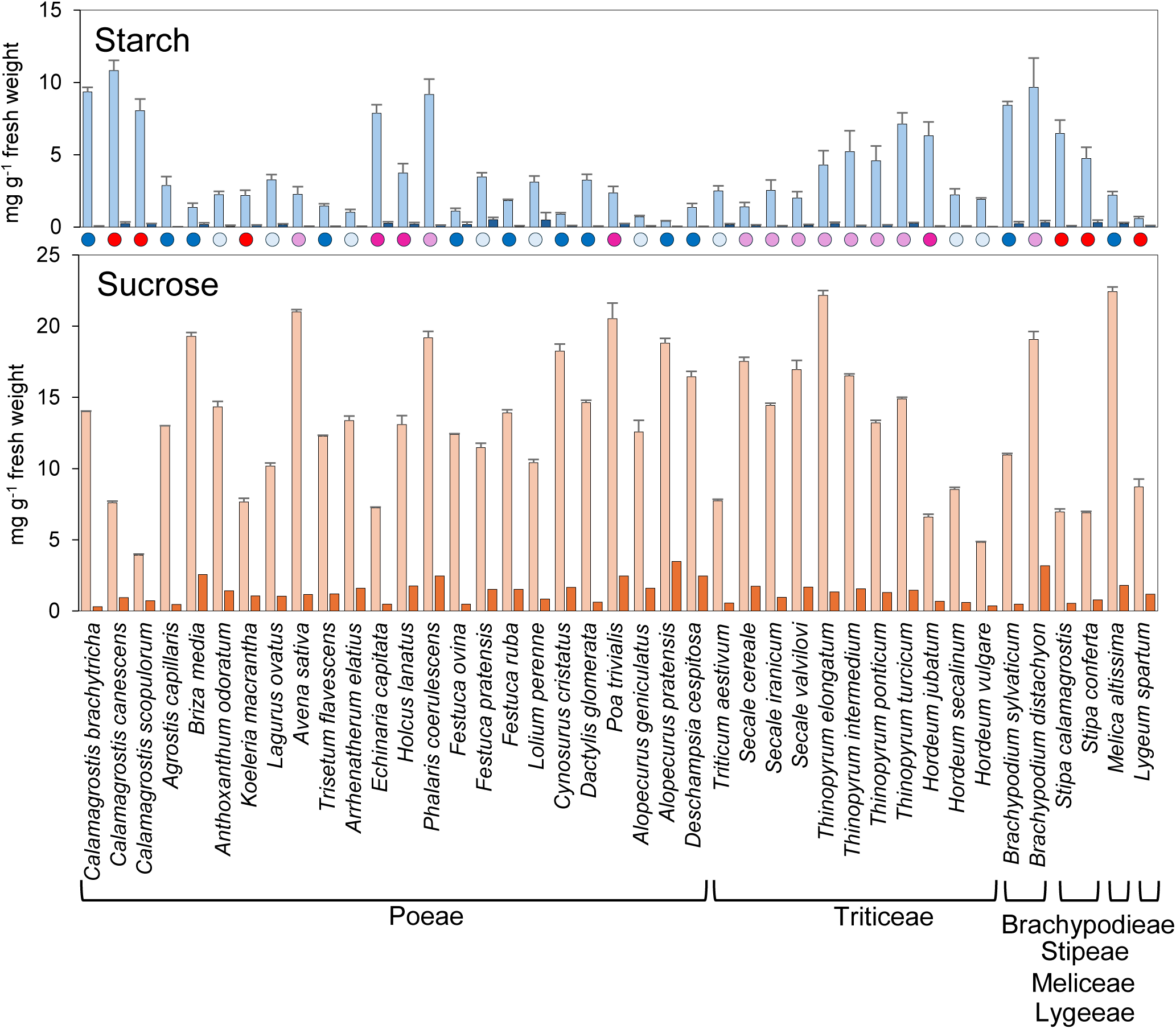
End-of-day and end-of night starch and sugar contents of grass leaves. Grasses were grown in a glasshouse with days of 10 h ambient light, 14 h dark, and ambient temperature. Sections 1 cm in length were taken from midway along the blade of the youngest fully expanded leaf of non-flowering plants. Species were grown and sampled in five batches, with harvests spread across the growing season between April and September. Each batch contained between four and eleven species, selected without phylogenetic bias. Measurements of starch (upper graph) and sucrose (lower graph) were made on insoluble and soluble extracts, respectively (see Methods). Coloured circles below the bars show dates of harvest in weeks before (-) and after (+) the summer solstice: -9 weeks 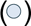, -6 weeks 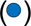, +6 weeks 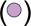, +9 weeks 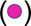, +16 weeks 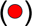. Tribes to which the species belong are shown below the species names. For each species, values are means of measurements on samples from five or six individual plants sampled at the end of the day, and from five or six different individual plants sampled at the end of the night. Bars show SE. For starch, pale blue bars are EoD values; dark blue bars are EoN values. For sucrose, pale orange bars are EoD values, dark orange bars are EoN values.

**Figure 2.**
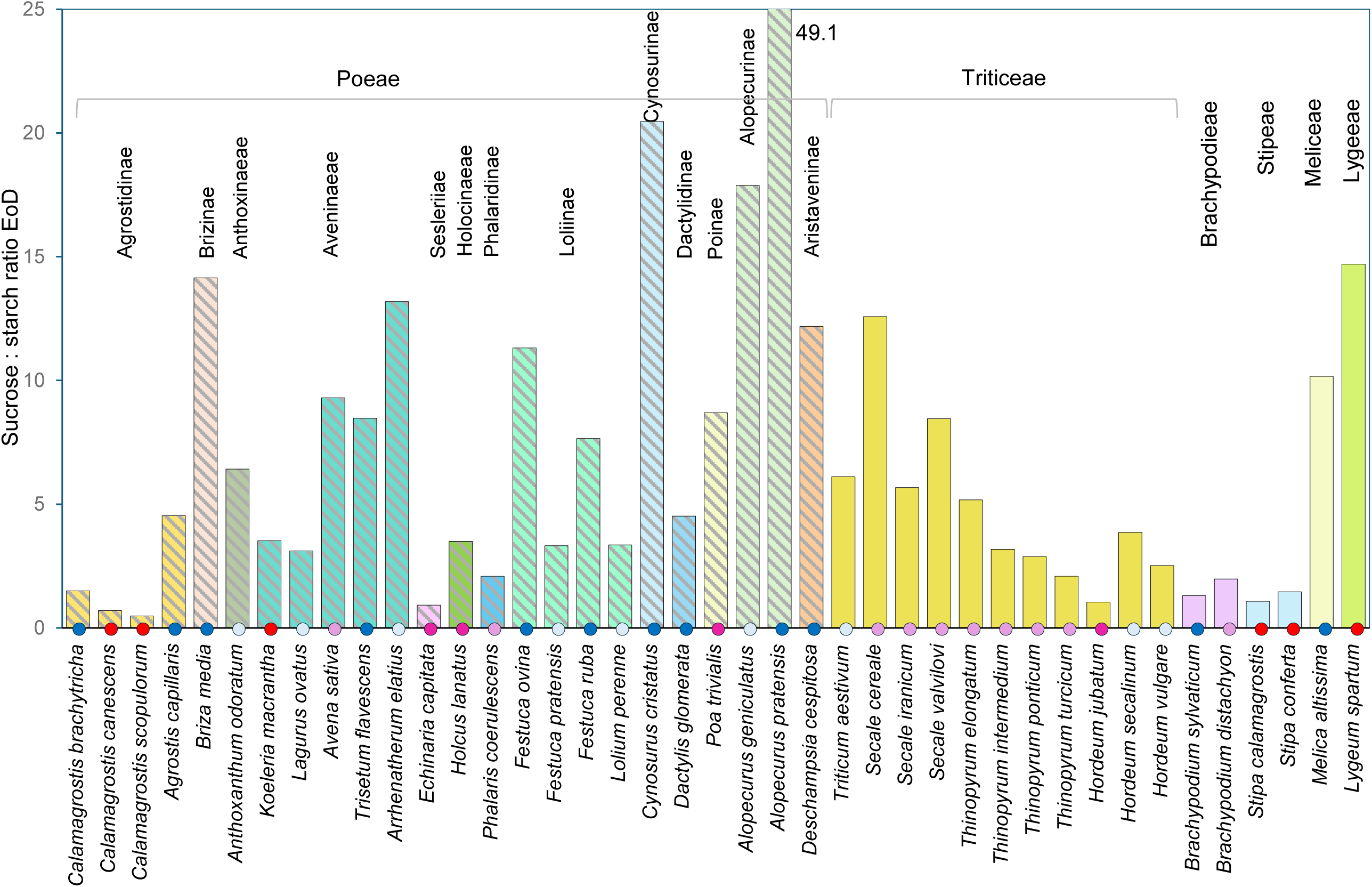
Ratio of sucrose to starch at the end of the day. Values were calculated from data in Figure 1. Hatched bars represent species in the tribe Poeae, coloured according to the subtribe of Poeae to which the species belongs (labelled at top). Solid bars represent species in five tribes other than Poeae (labelled at top). Coloured dots below the bars show the date on which plants were sampled, as for Figure 1. Note that the value for *Alopecurus pratensis* was 49.1.

All of the species accumulated both sucrose and starch during the 10-h light period (Figure 1). For all but three species, sucrose levels were higher than starch levels at the end of the day. In 32 out of the 41 species, the EoD sucrose:starch ratio was greater than 2, and in 10 of these species the ratio was greater than 10 (Figure 2). In nine species, the EoD sucrose:starch ratio was less than 2 and in three of these species it was between 0.5 and 1. The nine species with EoD sucrose:starch ratios of less than 2 included *Brachypodium* spp., S*tipa* spp. and *Calamagrostis* spp. There was no obvious relationship between the EoD sucrose:starch ratio and the tribe to which a given species belonged; in fact there was greater variation in EoD sucrose:starch ratios in species within the tribe Poeae than between species in the Poeae and species in other tribes in the Pooideae. For example, *Avena sativa*, *Trisetum flavescens* and *Phalaris coerulescens* from the tribe Poeae (all sampled with 6 weeks of the summer solstice) had EoD sucrose:starch ratios of 9.3, 8.5 and 2.1 respectively, a range that encompasses most of the species we sampled. In most cases species in the same genus had similar EoD sucrose:starch ratios. For example, the two species of *Alopecurus* in the study both had high ratios, and the two species of *Stipa*, two species of *Brachypodium* and three species of *Calamagrostis* all had markedly low EoD sucrose:starch ratios.

Both sucrose and starch were mobilised during the 14-h night (Figure 1, Figure S1). More than 90% of EoD sucrose was mobilised during the night in about half of the species; the other half of the species mobilised 80-90% of their EoD sucrose. Most of the EoD starch was also mobilised: all but five species mobilised more than 90% of their EoD starch and the remaining five species mobilised 80-90% of their EoD starch during the night (Figure S1). Overall, this initial analysis strongly indicated that 1) both sucrose and starch are subject to diel turnover in leaf blades of species in subfamily Pooideae; 2) in most species diel sucrose turnover is greater than starch turnover; 3) there are very substantial differences between species in the relative contributions of sucrose and starch to diel carbohydrate turnover; 4) differences between species in the relative contributions of sucrose and starch do not reflect their phylogenetic relationships at the level of tribes.

Two major considerations potentially affect interpretation of the data in Figures 1 and 2. First, for practical reasons, subsets of the 41 species were grown and sampled at intervals during spring and summer rather than as a single batch. As a consequence these subsets were subjected to different conditions of temperature, light intensity and light quality prior to sampling. Any or all of these factors might affect sucrose:starch ratios. Second, the presence of fructans in some species or conditions could potentially lead to over-estimation of sucrose contents. Commercially available invertases used in enzymatic sucrose assays may degrade fructo-oligosaccharides such as kestose as well as sucrose. The influence of these factors on interpretation of the data is discussed below.

### Influence of harvest date and fructans on measurements of leaf carbohydrate contents

It was apparent that the date of harvest had larger effects on average EoD sucrose contents than on average EoD starch contents (Figure 3). Although each harvest contained a different set of randomly-selected species, there was a strong trend in average EoD sucrose values across harvest dates. These values were highest in the two harvests six weeks either side of the solstice and lowest in the final harvest. By contrast, there was no obvious trend in average starch contents across all harvests. Average EoD sucrose values exceeded average EoD starch values in the first three harvests (at -9 weeks, -6 weeks and +6 weeks relative to the summer solstice), but fell within average EoD starch values by the final harvest. The differences in starch:sucrose ratios and absolute values between harvest dates cannot be due to differences in photoperiod since all plants were grown with 10 h light, 14 h dark: they are likely to be due to seasonal variation in average temperature, light levels and light quality.

**Figure 3.**
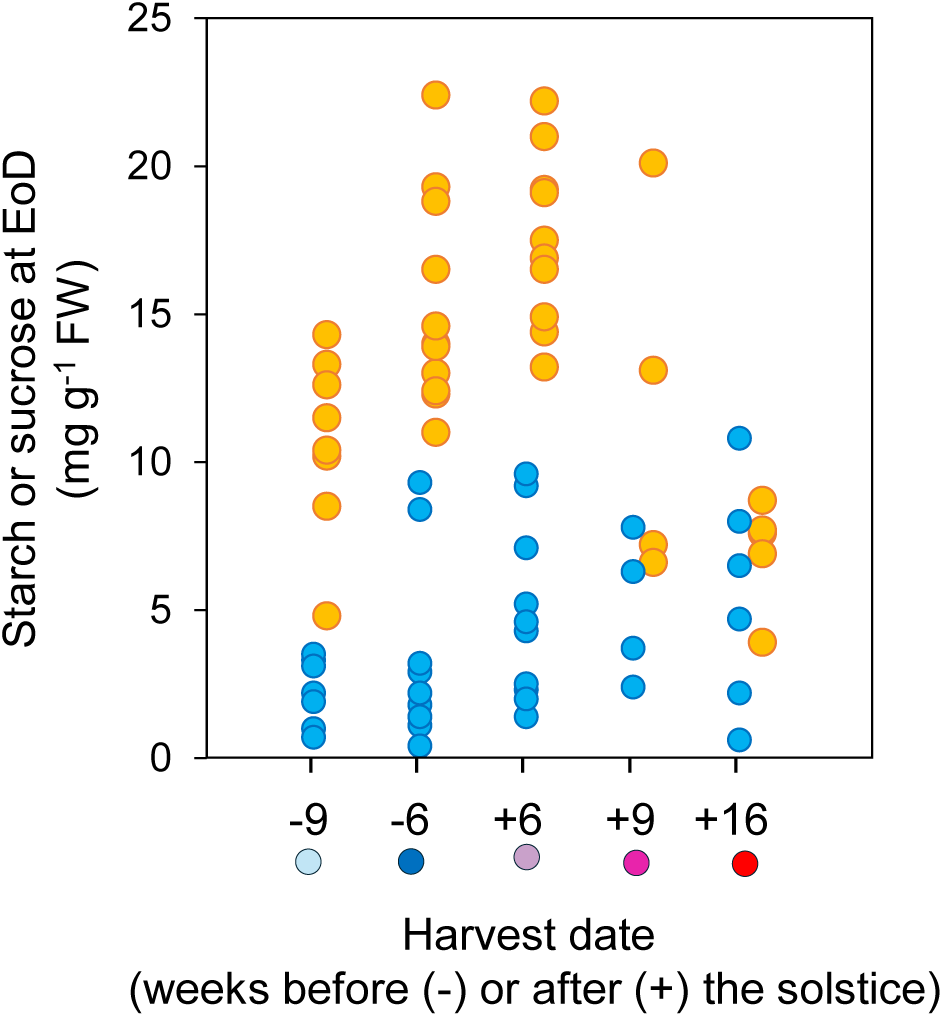
Impact of sampling date on starch and sugar contents of grass leaves. The graph shows the mean EoD starch and sucrose values for the species sampled on each of the five harvest dates (-9, -6, +6, +9, +16 weeks before/after the solstice). Values are taken from Figure 1. The coloured circles below the harvest dates correspond to those indicating harvest dates in Figures 1 and 2. Orange dots are sucrose values; blue dots are starch values. For clarity, sucrose values are slightly offset from starch values.

The marked effects of harvest date on sucrose accumulation did not affect the overall picture of carbohydrate turnover outlined in the previous section. Although the relative importance of sucrose and starch for diel carbohydrate turnover shifted through the season, examination of individual harvests continued to support the general idea that sucrose makes a contribution to diel carbohydrate turnover comparable with and in most species greater than that of starch. Data from species within individual harvests also supported the indications that, regardless of time of harvest, there was wide variation between species in the relative contributions of sucrose and starch to diel carbohydrate turnover (Figure 3). Within single harvests there were up to 3-fold differences between species in EoD sucrose contents, and up to 20-fold differences in starch contents.

We used enzymatic assays to assess whether potentially interfering fructans were present in extracts used for the starch and sucrose measurements shown in Figure 1. This analysis was done for 17 species including members of the tribes Poeae, Triticeae, Brachypodieae and Meliceae. For each extract, digestions with invertase, sucrase (an enzyme specific for sucrose) or fructanase yielded the same, essentially equimolar amounts of glucose and fructose, indicating that no fructo-oligosaccharides or larger fructans were present (Figures S2 and S3). Both invertase and fructanase are expected to yield an excess of fructose over glucose in the presence of small fructo-oligosaccharides, and fructanase is expected to have this effect in the presence of larger fructans. Digestion of standard samples of sucrose and the fructans levan and inulin with fructanase yielded the ratios of glucose to fructose expected from the manufacturer’s specifications (not shown), in both the absence and the presence of leaf extract (Figure S2C), showing that the apparent absence of fructans was not due to inhibition of fructanase by components of leaf extracts.

As a further check that the apparent absence of fructans in leaves grown in 10 h light, 14 h dark was not artefactual, we examined yields of hexoses in extracts of leaves of 10 species or cultivars grown in long days and high light (18 h light at 400 μmol quanta photosynthetically active radiation (PAR) m^-2^ s^-1^, 6 h dark) in a controlled environment room (CER). Long-day conditions are reported to promote fructan accumulation in several of the species in our survey (see Introduction). Our results showed that fructans did indeed accumulate in these conditions. Yields of hexose from digestions with invertase, invertase+sucrase or invertase+fructanase revealed the presence of compounds in addition to sucrose that were susceptible to invertase hydrolysis (putatively fructo-oligosaccharides), and of larger fructans (Figure S3 and see Methods).

### Diel turnover of carbohydrate and its response to light intensity and day length differs between species under controlled growth conditions

To provide further information on the control of starch and sucrose metabolism in grass leaves, we investigated the impact of light intensity and day length on the extent and pattern of diel turnover of these carbohydrates. We chose six species that spanned the range of variation present in the glasshouse experiment shown in Figures 2 and 3, and grew them in a CER. The species were *Brachypodium distachyon* and *Phalaris coerulescens* with markedly low EoD sucrose:starch ratios, *Avena sativa* and *Melica altissima* with high EoD sucrose:starch ratios, and *Triticum aestivum* and *Thinopyrum turcicum* with intermediate EoD sucrose:starch ratios. Plants were grown in combinations of short or long days (respectively 8 h light, 16 h dark; and 16 h light, 8 h dark) and low, intermediate or high light (respectively 100, 200 and 400 μmol quanta PAR m^-2^ s^-1^). As for the glasshouse experiment, measurements were made at the end of the day and the end of the night on sections taken from midway along the blades of youngest fully expanded leaves, from six different plants. Figure 4A shows mean values of starch and sucrose for each species at EoD and EoN at low, intermediate and high light intensities, under short and long days. The differences in EoD sucrose:starch ratios observed between these species in the glasshouse were largely reproduced under both short- and long-day conditions in the CER. *Brachypodium* had low EoD sucrose:starch ratios, *Phalaris, Triticum* and *Thinopyrum* had intermediate ratios, and *Melica* and *Avena* had high ratios (Figure 4B). Responses to day length and light levels, described below, showed some general trends but also differed markedly between species.

**Figure 4.**
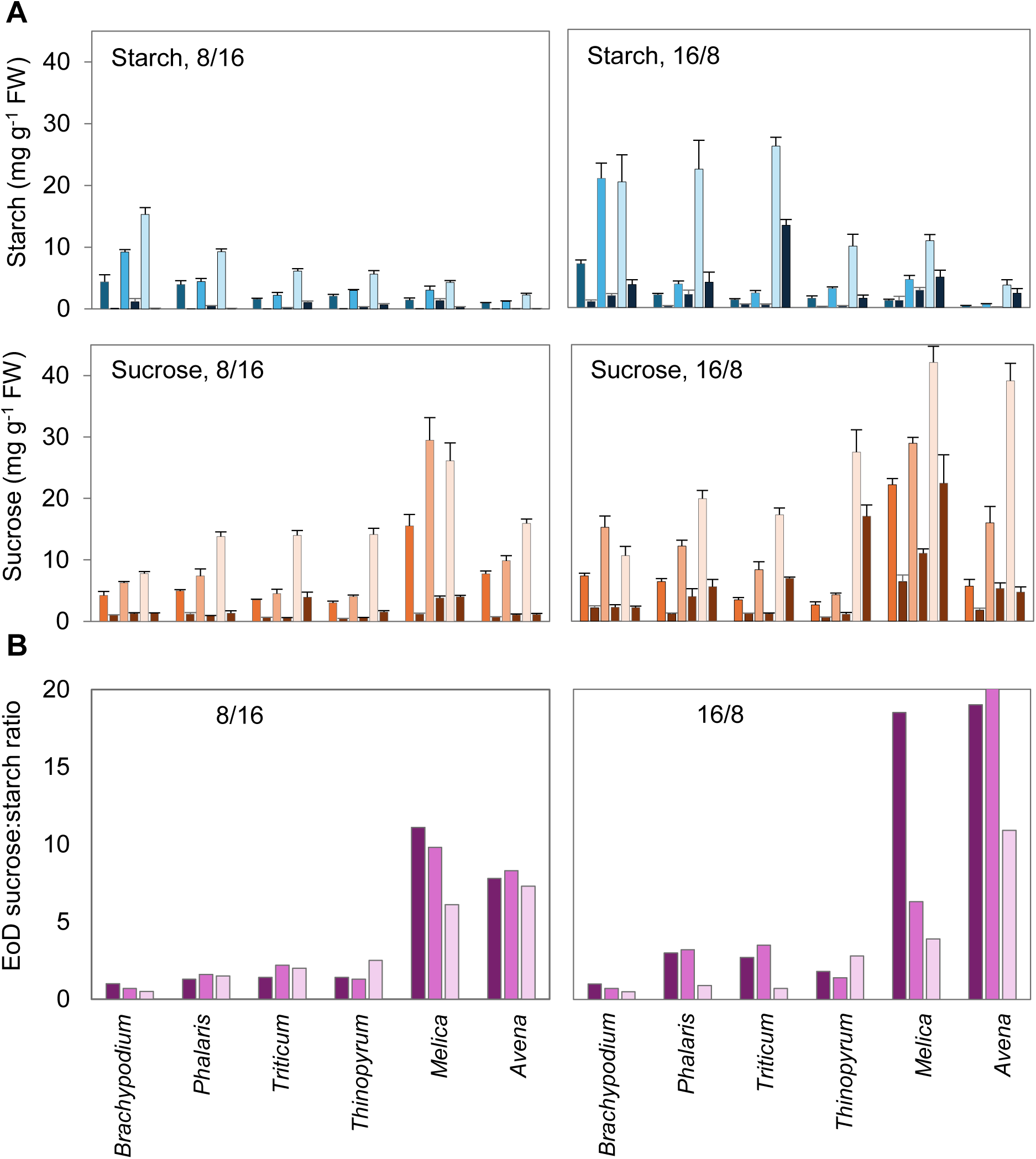
Impact of day length and light intensity on starch and sucrose contents in leaves of six grass species. **A** The graphs show results for plants grown in short day (left, 8/16) and long day (right, 16/8) conditions in a CER. Upper graphs show starch contents for plants grown with 100, 200 or 400 μmol quanta PAR m^-2^ s^-1^, at EoD (blue-grey, mid-blue and pale blue bars respectively) and EoN (black bars). Lower graphs show sucrose contents for plants grown with 100, 200 or 400 μmol quanta PAR m^-2^ s^-1^, at EoD (dark orange, mid-orange and pale orange respectively) and EoN (brown bars). For each harvest point, values are means of measurements on samples from six individual plants sampled at EoD, and from six different plants sampled at EoN. Bars show SE. Note that all y axes are on the same scale to facilitate comparisons across species, light intensities and day lengths. **B** Sucrose:starch ratios at EoD calculated from **A**. Left: short-day conditions (8/16). Right: long-day conditions (16/8). Dark purple, mid-purple and pale purple bars represent plants grown with 100, 200 or 400 μmol quanta PAR m^-2^ s^-1^ respectively.

In general, plants grown in long days contained more sucrose and more starch at EoD than plants grown in short days, but the difference between short- and long-day values was mostly less than two-fold. The impact of light intensity on the EoD sucrose:starch ratio was relatively small (two-fold or less in most cases) and differed with day length and between species (Figure 4B). *Brachypodium* and *Melica* plants contained more starch relative to sucrose at high than at lower light levels in both short and long days, and this was also true of *Phalaris*, *Triticum* and *Avena* in long days.

In short days, sucrose and starch depletion at night resembled that in the glasshouse experiment, in that in most instances 20% or less of both carbohydrates remained at the end of the night, regardless of daytime light intensity (Figure 5A). In long days, less of the EoD sucrose and starch was depleted at night. More than 20%, and in some instances more than 40%, of EoD sucrose and/or starch remained at the end of the night. However, despite this substantial difference in EoN sucrose and starch levels between long- and short-day leaves, for a given species the absolute amounts of sucrose, starch and total carbohydrate (sucrose+starch) lost during the night were rather similar in long and short days (Figure 5B, Figure S4).

**Figure 5.**
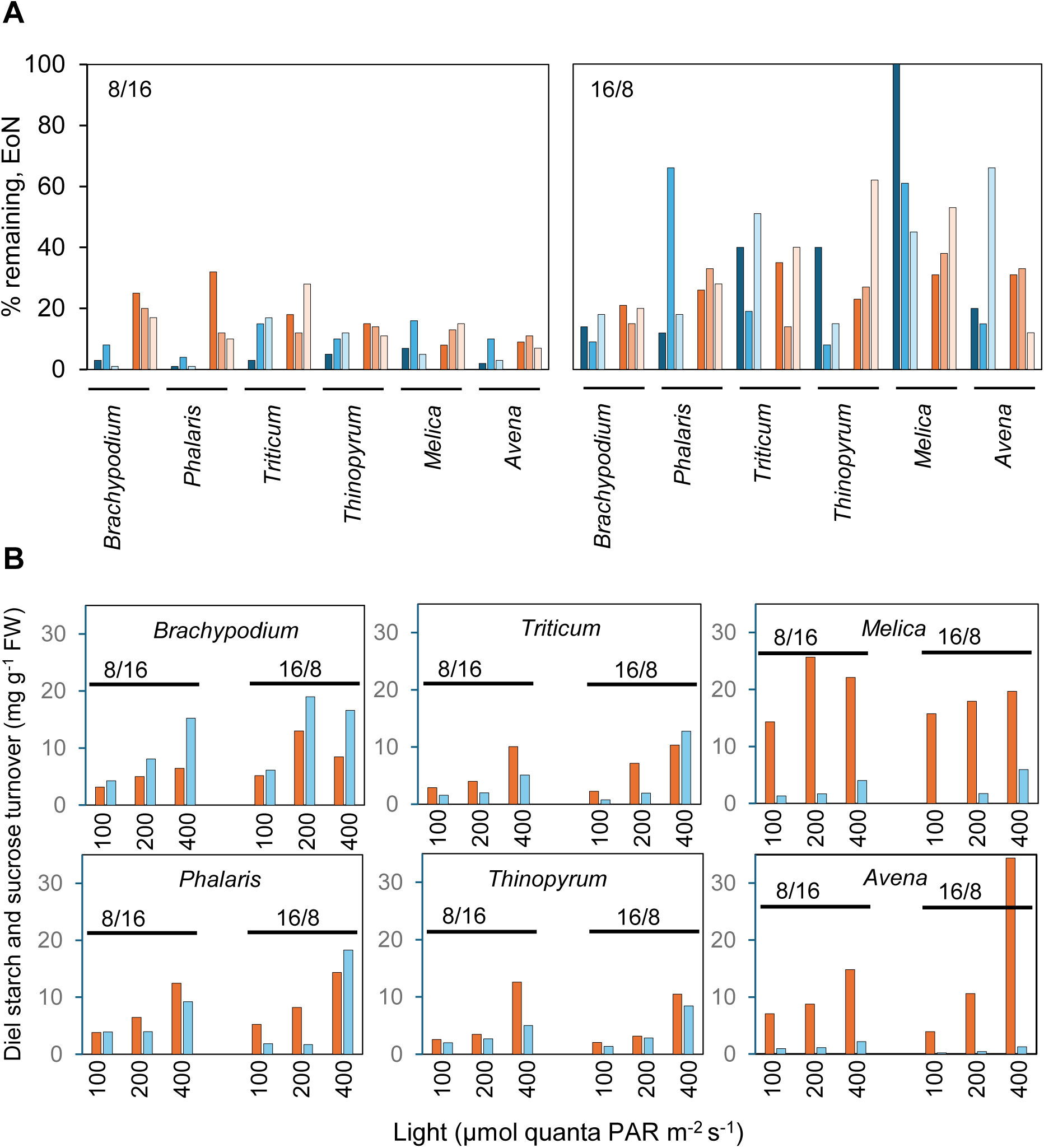
Effect of day length and light intensity on diel turnover of sucrose and starch in leaves of six grass species. **A** Percentage of EoD starch (blue bars) and sucrose (orange bars) remaining at EoN in short (8/16, left graph) and long (16/8, right graph) days, calculated from the data in Figure 4A. Dark, mid- and pale coloured bars represent plants grown at 100, 200 and 400 μmol quanta PAR m^-2^ s^-1^, respectively. **B** Amounts of starch (blue bars) and sucrose (orange bars) turned over during the night in leaves of six grass species grown under short days (left side of each graph) or long days (right side of each graph), calculated from the data in Figure 4A. Dark, mid- and pale coloured bars represent plants grown at 100, 200 and 400 μmol quanta PAR m^-2^ s^-1^, respectively.

### Fructans do not appear to undergo diel turnover in *Avena* and *Triticum*

The presence of fructans in leaf extracts could affect accuracy and interpretation of our data in two ways. First, as described above, hydrolysis of fructans could compromise the accuracy of sucrose assays. Second, fructans might undergo diel turnover. If this were the case, measurements of starch and sucrose alone would not provide a complete picture of the diel control of carbohydrate availability in grass leaves.

We showed above that fructans were present under high light and long day conditions in leaves of several species in our study (Figure S2). To look for diel turnover, we used Dionex HPLC (HPAEC) analysis of leaf extracts to investigate the soluble carbohydrate content of *Triticum* and *Avena* leaves at the end of the day and the end of the night. This method revealed the presence in high-light, long-day-grown leaves of compounds with elution times greater than those of sucrose and hexoses and similar to or greater than those of standard fructo-oligosaccharides. These compounds were absent from leaves of low-light, 12/12- grown material (examples of HPAEC traces are in Figure S5). They are thus likely to be fructans of a range of different sizes. For both *Triticum* and *Avena* leaves grown in high light and long days, the areas under the putative fructan profiles at EoN and EoD (normalised on a fresh weight basis and expressed in arbitrary units g^-1^ fresh weight) were not statistically significantly different (Student’s T-test P>0.05, Figure 6). This method does not allow quantitative comparison of sucrose and fructan levels, but the apparent absence of diel fluctuation of fructans under high light and long days provides assurance that fructan turnover is unlikely to lead to over-estimation of sucrose turnover in these conditions. However it remains possible that the high EoN levels of sucrose seen in high light and long days in some species (Figure 5) are overestimates due to some fructan hydrolysis.

**Figure 6.**
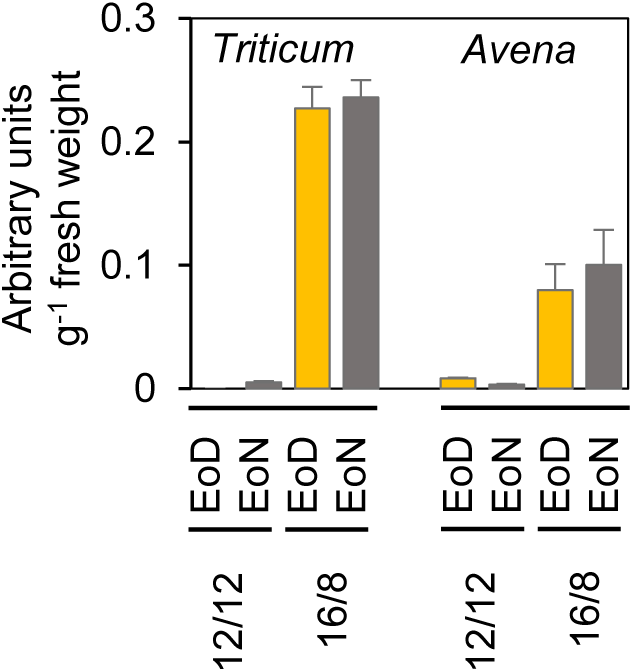
Effect of day length and light intensity on EoD and EoN fructan contents in *Triticum* and *Avena* leaves. Fructan contents at EoD (yellow bars) and EoN (grey bars) of the youngest fully expanded leaves of *Triticum* (left) and *Avena* (right). Plants were grown in either 12-h, low light days (100 μmol quanta PAR m^-2^ s^-1^) or long, high light days (16/8, 400 μmol quanta PAR m^-2^ s^-1^). Extractions were as for starch and sucrose measurements. The soluble fraction was passed through anion and cation exchangers then applied to a Carbopac PA1 column on a Dionex HPLC and eluted with a gradient of sodium hydroxide and sodium acetate. Figure S5 shows evidence of linearity of detector responses, typical elution profiles of standards and samples, and region of the sample profiles used for measurement. Values are in arbitrary units per g fresh weight, normalised to an internal standard. They are means ±SE of measurements on extracts of 6 different plants.

### Rates of sucrose and starch depletion during the night are not linear

Results thus far showed that the relative amounts of sucrose and starch lost from leaf blades during the night differed profoundly between species. To investigate whether the patterns of sucrose and starch depletion also differed between species, we measured sucrose and starch contents of leaves at intervals of 2 h over 24 h in a CER with 12 h light, 12 h dark and 400 μmol quanta PAR m^-2^ s^-1^). In these conditions carbohydrate turnover was high and little remained at the end of the night in almost all species (Figure 7), permitting accurate measurements and minimising difficulties of interpretation presented by differences between species in the extent of turnover. These measurements were made on two species that had low EoD sucrose:starch ratios in previous experiments (*Brachypodium* and *Phalaris*), four that had “intermediate” ratios (*Lagurus, Thinopyrum turcicum* and *Echinaria*) and three that had high EoD sucrose:starch ratios (*Avena*, *Melica* and *Alopecurus geniculatus*).

**Figure 7.**
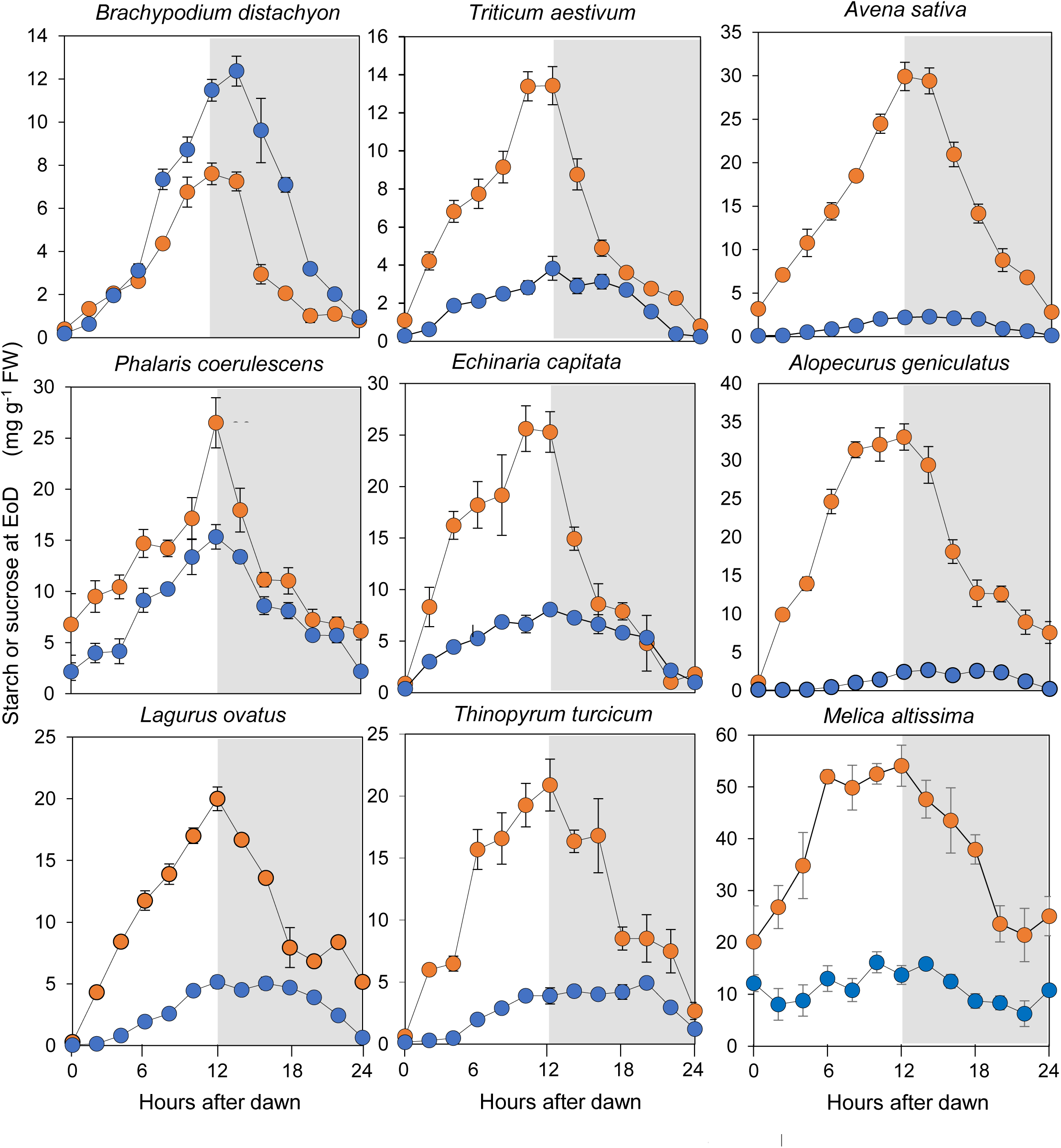
Sucrose and starch contents over a diel cycle for nine grass species. Plants were grown in a CER with 12 h light, 12 h dark and 400 μmol quanta PAR m^-2^ s^-1^ and sampled at 2-h intervals for sucrose (orange) and starch (blue) measurements. Values are means ±SE of measurements on extracts of 6 different plants. Where error bars are not visible, they are smaller than the symbols. Different plants were sampled at each time point.

There was good correspondence of EoD sucrose:starch ratios between this 24-h time-course experiment (Figure 7) and the glasshouse (Figure 2) and daylength/light intensity (Figure 4B) experiments. As previously, *Brachypodium* and *Phalaris* had low/intermediate EoD sucrose:starch ratios, *Triticum*, *Lagurus, Thinopyrum* and *Echinaria* had intermediate ratios, and *Avena*, *Melica* and *Alopecurus* had particularly high EoD sucrose:starch ratios. Unlike the glasshouse and daylength/light intensity experiments, in the 24-h time-course experiment *Melica* leaves retained substantial amounts of starch and sucrose at EoN and there was no obvious diel fluctuation of starch content (Figure 7).

The pattern of depletion of both sucrose and starch during the night (Figure 7) appeared non-linear. Normalisation of data to EoD values indicated that in all species except *Melica* sucrose mobilisation occurred earlier and faster than starch mobilisation, then slowed, whereas starch mobilisation was initially slow then accelerated in the later part of the night (Figure 8). This difference was reflected in the percentage of sucrose versus starch mobilised in the first half of the night. In all nine species about 60 to 80% of EoD sucrose was mobilised in the first half of the night (Table 1). By contrast, 32% or less of EoD starch was mobilised before the middle of the night in seven out of the nine species, four of which mobilised less than 10% of EoD starch over this period. The two species with the lowest EoD sucrose:starch ratios, *Brachypodium* and *Phalaris*, mobilised similar amounts of their EoD starch in the first half and the second half of the night (41% and 55% mobilised by ZT18 in *Brachypodium* and *Phalaris* respectively). (Note that *Melica* was excluded from further analysis of starch mobilisation because there were no clear trends in the data for this species.)

**Figure 8.**
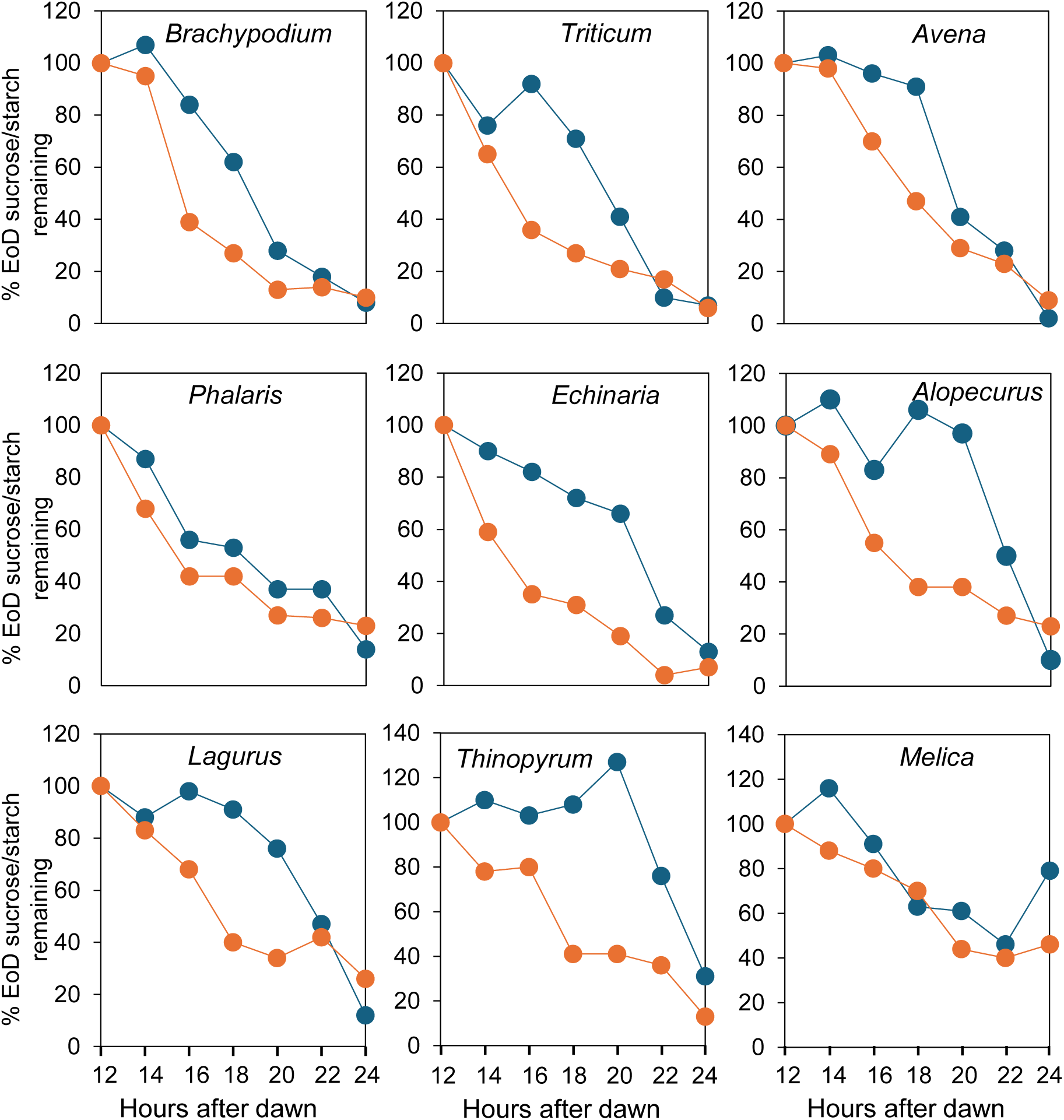
Sucrose and starch contents over the night, normalised to the respective EoD values. Data for the 12-h period EoD to EoN (dark period) in the experiment shown in Figure 7 are normalised to EoD values to facilitate comparison of the patterns of depletion of sucrose (orange) and starch (blue).

**Table 1.** Sucrose and starch depletion in the first half of the night. Data are starch or sucrose contents halfway through the night (ZT18), expressed as a percentage of the total loss between ZT12 and ZT24 : 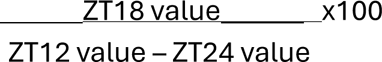 Values are taken from Figure 7. For *Melica*, starch depletion showed no clear trend over the night.

| Species | Sucrose at 18 h as % of loss between 12 and 24 h | Starch at 18 h as % of loss between 12 and 24 h |
|---|---|---|
| *Brachypodium* | 81 | 41 |
| *Phalaris* | 76 | 55 |
| *Lagurus* | 81 | 10 |
| *Triticum* | 78 | 29 |
| *Echinaria* | 74 | 32 |
| *Thinopyrum* | 68 | 0 |
| *Avena* | 58 | 9 |
| *Alopecurus* | 80 | 0 |
| *Melica* | 56 | - |

The results above suggest that control of starch mobilisation in pooid grass species differs from that in Arabidopsis, in which the rate of mobilisation during the night is both linear and sufficient to exhaust most of the EoD starch by dawn (Graf et al., 2010; Scialdone et al., 2013). Although most of the EoD starch was exhausted by dawn in all of the grasses in the 24-h time-course experiment, starch mobilisation appeared biphasic in species other than *Brachypodium* and *Phalaris*: it was faster in the second than in the first half of the night (Table 1). Mobilisation in the second half of the night (ZT18-24) approximated to linearity (Figure 8; R^2^ ∼0.9) or greater for all but *Thinopyrum*, for which R^2^ was 0.76 for ZT18-24, and 0.99 for ZT20-24. Extrapolation of the linear rate over this period revealed that, in all species, it would exhaust the starch present at ZT18 by a point close to dawn (Figure 9: between ZT23.9 and ZT25.3 for six species, ZT27.1 for the remaining two).

**Figure 9.**
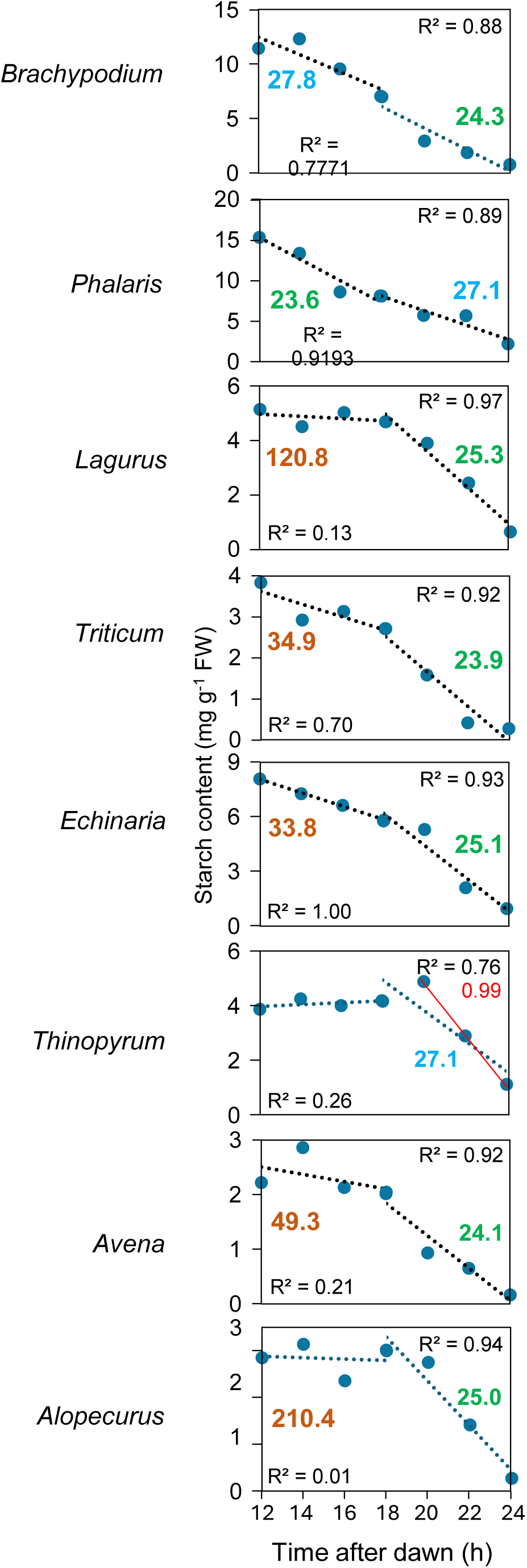
Pattern of starch depletion over the night. Data for starch for the 12-h period EoD to EoN in the experiment shown in Figure 7 with lines of best fit (dotted) plotted separately for ZT12 to ZT18 and ZT18 to ZT24. R^2^ values are provided at the lower left and upper right of each graph for the ZT12-18 and ZT18-24 lines respectively. Coloured numbers on each graph show the time after dawn (ZT0) at which starch content would reach zero based on extrapolation of the line of best fit. Left, time for the ZT12-18 line; right, time for the ZT18-24 line. Green numbers are values within 1.3 h of ZT24; blue numbers are values within 3.8 h of ZT24; brown numbers are values differing from ZT24 by more than 9 h. No value is given for *Thinopyrum* from ZT12 to ZT18 because the slope of the line of best fit was positive.

By contrast, the rate of starch consumption in the first half of the night (ZT12-ZT18) was too slow to exhaust starch reserves around ZT24 in all species except *Brachypodium* and *Phalaris* (Figure 9). Extrapolation of a straight line fitted to data from this period gave theoretical times for exhaustion of starch reserves ranging from about ZT34 for *Triticum* and *Echinaria,* ZT49 for *Avena* and ZT120 and ZT210 respectively for *Lagurus* and *Alopecurus*. For *Brachypodium* and *Phalaris*, the two species with the lowest EoD sucrose:starch ratios, the rates of starch consumption in the first and second halves of the night were similar. Both rates approximated to linearity, and would theoretically exhaust starch reserves within 4 h of ZT24.

In summary, in eight species of pooid grasses starch depletion in the second half of the night (ZT18-24) was linear and at a rate that would exhaust the starch present at ZT18 by a point close to dawn. For six out of the eight species, the rate of starch loss in the first half of the night was slower than in the second half, and too slow to exhaust EoD starch reserves by dawn. For the two species with the lowest sucrose to starch ratios, starch loss was essentially linear throughout the night, and was at a rate that would exhaust EoD starch reserves close to dawn.

## DISCUSSION

### Sucrose makes a major contribution to night-time carbohydrate availability in grass leaves

Our results greatly extend knowledge of diel carbohydrate turnover in leaves of pooid grasses. Despite very substantial variation between species in diel patterns of CHO turnover, it is clear that sucrose rather than starch is the major contributor to carbohydrate availability at night in the family as a whole. The species we examined covered a broad spectrum of the Pooideae, and in all cases sucrose depletion at night exceeded or was similar in magnitude to the consumption of starch. Both the dominance of sucrose and the substantial variation between species in diel patterns of carbohydrate turnover were robust features of our data, maintained over a wide range of constant and variable growth conditions. We found no evidence that fructans made a significant contribution to diel carbohydrate turnover in our study. However it remains possible that diel fructan turnover is significant in other species and growth conditions.

Leaves of other pooid grasses are also reported to mobilise more sucrose than starch during the night (e.g. *Festuca arundinacea*, Shewmaker et al., 2006; *Phleum pratense*, Bertrand et al., 2008; *Poa* spp., Borland and Farrar, 1985), or to have very low starch contents (Sakai and Hayashi, 1973). Surveys of the distribution among angiosperms as a whole of sucrose-dominated versus starch-dominated leaf carbohydrate content have reported very wide variation between species (e.g. Chatterton et al., 1989; Goldschmidt and Huber, 1992; Barbehenn et al., 2004; Asao et al., 2025). There are numerous measurements of starch and sucrose levels at single timepoints in leaves of many species, but such measurements do not reveal the relative importance of starch and sucrose for carbohydrate availability at night. Species in which both sucrose and starch were shown to undergo diel fluctuation include spinach (*Spinacia oleracea*; Gerhardt et al., 1987), sugar beet (*Beta vulgaris*; Fondy and Geiger, 1982), tomato (*Lycopersicum esculentum*; Matsuda et al., 2014), soybean (*Glycine max*; Chatterton and Silvius, 1979; Hewitt et al., 1985), maize (*Zea mays*; Kalt-Torres et al., 1987; Czedik-Eysenberg et al., 2016; Liang et al., 2019, 2021), sugar cane (*Saccharum officinarum*; Du et al., 2000; Da Silva et al., 2023), white clover (*Trifolium repens*; Gordon et al., 1987), *Lotus japonicus* (Vriet et al., 2010), grape vine (*Vitis vinifera*; Dayer et al., 2020), red oak (*Quercus rubra*; Gersony et al., 2020), field bean (*Vicia faba*; Fondy et al., 1989; Wacker et al., 2025), almond (*Prunus dulcis*; Tixier et al., 2018) and sunflower (*Helianthus annuus*; Wacker et al., 2025). Wide variation in ratios of sucrose to starch turnover were reported across these studies. We discuss the possible origins of the diversity of sucrose to starch turnover within the Pooideae in more detail below.

### The rate of sucrose depletion at night may be related to sucrose content

The amount of sucrose that accumulated during the day in pooid grass leaves was dependent on both daylength and light intensity, in a species-specific manner. Except in high light and long days, most (75-80% or more) of EoD sucrose was mobilised during the night. The rate of sucrose depletion during the night appeared non-linear (near-exponential) for several species (Table 1; Figure 8), including species with both very high and very low EoD sucrose:starch ratios. However, the data overall were not adequate to allow generalisations about linear versus exponential rates of sucrose depletion (Figure S6). Exponential depletion would imply that the process may be controlled largely by sucrose content alone.

Non-linear depletion of sucrose in leaves at night has been reported for a wide range of monocot and eudicot species. In barley, sucrose depletion at night was near-exponential in plants grown under a wide range of different light intensities and day lengths (Gordon et al., 1982; Sicher et al., 1984; Farrar and Farrar, 1985; Müller et al., 2018; Liang et al., 2021). This was the case not only under constant conditions but also when daylength was unexpectedly increased from 12 h to 16 h or 20 h: the close relationship between EoD sucrose level and the rate of sucrose depletion was retained even though the EoD sucrose level was higher under the longer day (Müller et al., 2018). Near-exponential depletion of sucrose was also observed in *Poa* spp. (Borland and Farrar, 1985), wheat (Balaguer et al., 1995; Annunziata et al., 2026), sugar beet (Fondy and Geiger, 1982), sugar cane (Du et al., 2000) and spinach (Gerhardt et al., 1987). Based on kinetic analyses of metabolite pools in leaves labelled with ^14^CO_2_, it has been suggested that sucrose depletion is a two-phase process, consisting of an initial, rapid phase in which a cytosolic pool of sucrose is exported from the leaf, overlapping with a second, slower phase in which vacuolar sucrose moves via tonoplast sucrose transporters to the cytosol for export (Sicher et al., 1984; Gordon et al., 1980a, 1980b, 1982; Geiger et al., 1983; Farrar and Farrar, 1985; Lattanzini et al., 2012). Consistent with these suggestions, loss of the rice tonoplast sucrose transporter SUT2 reduces sucrose depletion in leaves at night and changes the pattern of sucrose depletion from exponential to linear (Eom et al., 2011; Müller et al., 2018).

Additional intrinsic factors are also likely to influence the kinetics of sucrose depletion in the leaf at night. For export via the phloem, sucrose from a mesophyll cell must reach the phloem parenchyma, where it is exported to the apoplast via SWEET transporters for uptake and phloem loading by phloem companion cells (Braun, 2022). The route to the phloem parenchyma involves symplastic movement via plasmodesmata through several other cells: perhaps other mesophyll cells, and then bundle sheath cells (consisting of outer parenchymatous sheath cells and inner mestome sheath cells in pooid grasses: Williams et al., 1989; Leegood, 2008; Braun, 2022). Movement along this route will depend on the maintenance of a sucrose concentration gradient between the mesophyll and the phloem parenchyma. As sucrose concentration in the mesophyll falls at night, the steepness of the gradient will presumably also fall, slowing the rate at which sucrose in the mesophyll cell is depleted as the night progresses. The kinetics of depletion of sucrose during the night may also be influenced by magnitude and connectivity of pools of sucrose in the cytosol and vacuoles of the bundle sheath and phloem-associated cells, and changes in the requirement for sucrose consumption in companion-cell respiration to support phloem loading. The rate of conversion of starch to sucrose in mesophyll cells during the night will also affect the kinetics of sucrose depletion. Our data suggest that the contribution of starch mobilisation to the sucrose pools of mesophyll cells will be significant throughout the night in species such as *Brachypodium* and *Phalaris* with low EoD sucrose:starch ratios and relatively constant rates of starch mobilisation, but of significance only in the latter part of the night in species with high EoD sucrose:starch ratios and very slow starch mobilisation in the first part of the night.

### Starch consumption is non-linear when end-of-day sucrose:starch ratios are high

The pattern of starch consumption during the night differed between grass species, and was clearly regulated. In *Brachypodium* and *Phalaris*, the two species with the lowest EoD sucrose:starch ratios in the time-course experiment (0.7 and 1.7 respectively), consumption in plants grown in 12 h high light, 12 h dark was essentially linear, and was such that starch was exhausted approximately at dawn. This is consistent with the idea that degradation is controlled by a mechanism involving the circadian clock, as is the case in Arabidopsis (Graf et al., 2010; Scialdone et al., 2013; Ishihara et al., 2022). In *Lagurus*, *Triticum*, *Echinaria*, *Thinopyrum, Avena* and *Alopecurus*, in which EoD sucrose:starch ratios were between 3 and 15, starch consumption was very slow during the first part of the night then accelerated later in the night. The linear rate for the first part of the night was too slow to result in exhaustion of starch by dawn: projected times of starch exhaustion at these rates were between 10 h and 210 h after dawn. By contrast, starch consumption in the latter part of the night was linear and at a rate projected to exhaust starch within 1 to 3 h of dawn.

The patterns of night-time carbohydrate depletion in our study of eight pooid grasses are strikingly similar to those reported for the pooid grass barley (*Hordeum vulgare*). In overview, Gordon and colleagues showed that starch consumption was linear when the EoD sucrose:starch ratio was low (due to growth in low light) but was biphasic, with a slow initial rate and an acceleration in the latter part of the night, when EoD sucrose:starch ratios were high (due to growth in high light; Gordon et al., 1980a). Importantly, as in our study, the rate in the second half of the night appeared to be adjusted to the starch content roughly halfway through the night, such that it was adequate to permit near-complete exhaustion of starch by dawn regardless of the EoD sucrose:starch ratio. Gordon and colleagues speculated that in the first part of the night starch degradation was inhibited by high levels of sucrose, and that starch degradation accelerated once sucrose levels had fallen to a particular threshold. A lag in the onset of starch consumption at night in barley leaves was also reported by Sicher et al., 1984) and Müller et al. (2018), and lags have been reported for other species, for example sugar beet (Geiger and Batey, 1967; Fondy and Geiger 1982).

Recent research in Arabidopsis shows that starch degradation in leaves may indeed be inhibited by sucrose, and provides a mechanistic explanation for this phenomenon apparently applicable to a wide range of angiosperm species including both monocots and eudicots (Fitchner and Lunn, 2021; Ishihara et al., 2022; Annunziata et al., 2026). In brief, starch degradation over the diel cycle is under the control of a mechanism that continuously divides starch content by time remaining until dawn (measured by the circadian clock) and adjusts degradation to a rate that will exhaust starch supplies around dawn (Scialdone et al., 2013; Ishihara et al., 2022), maximising the efficiency of use of assimilated carbon over the diel cycle in carbon-limited conditions (Sulpice et al., 2014). The rate of degradation set by this control mechanism (the arithmetic division mechanism) can be suppressed by sucrose, via the sucrose-signalling metabolite trehalose 6-phosphate (Tre6P). When sucrose levels are high, Tre6P mediates feedback inhibition of starch degradation (Martins et al., 2013; dos Anjos et al., 2018; Ishihara et al., 2022). The presence of the Tre6P-synthesising enzyme TPS1 primarily in cells associated with the vasculature has led to the proposal that Tre6P levels directly reflect the size of a sucrose export pool, and that it diffuses symplastically from the vasculature into starch-degrading mesophyll cells (Fichtner et al., 2020; Fichtner and Lunn, 2021; Tonetti et al., 2025; Annunziata et al., 2026). Thus high levels of sucrose export can override the arithmetic division mechanism to give a lower rate of starch degradation than would permit exhaustion of starch reserves by dawn.

Based on these new insights, we suggest that the high sucrose contents and rates of export in leaves in the first part of the night in six out of our eight grass species, and in barley (Gordon et al., 1980a, Sicher et al., 1984, Müller et al., 2018), result in levels of Tre6P high enough to suppress the rate of starch degradation, such that it is insufficient to exhaust starch by dawn. The depletion of sucrose in the first part of the night then lowers Tre6P levels to a point at which Tre6P-mediated suppression of starch degradation is relieved, allowing the arithmetic division mechanism to set a rate sufficient to exhaust starch around dawn. In the two grass species with the lowest levels of sucrose relative to starch in the first part of the night (*Brachypodium* and *Phalaris*) and in the low-light-grown barley plants of Gordon and colleagues, we speculate that Tre6P does not reach levels sufficient to override the arithmetic control mechanism, giving a linear rate of starch degradation throughout the night that exhausts starch around dawn. Our suggestions are strongly supported by recent evidence that Tre6P levels are correlated with sucrose levels in wheat leaves, and remain high in the first part of the night rather than falling immediately at the onset of darkness as is the case in Arabidopsis leaves (Annunziata et al., 2026). Thus the arithmetic division mechanism plus its capacity for suppression by sucrose via Tre6P may represent an underpinning mechanism for control of carbohydrate availability that operates in angiosperms as a whole.

The arguments above suggest that the difference in diel carbohydrate turnover in mature leaves between the pooid grasses, in which both sucrose and starch contribute, and Arabidopsis, in which only starch contributes, may not represent a fundamental distinction between the Pooideae and eudicots. Although much more starch than sucrose is turned over in leaves of Arabidopsis and its close relative oilseed rape (*Brassica napus*; King et al., 1997; Mitchell et al., 2020; Lu et al., 2022), there are many reports of significant turnover of both sucrose and starch in leaves of diverse eudicots (see above). However, we are not aware of eudicots in which almost all of the diel turnover of carbohydrates in leaves is as sucrose, as is the case in *Avena* and *Melica* in our study and in many other monocots (e.g. *Allium* species; Parkin, 1899; Chapman, 1924; Pollock, 1986).

We emphasise that the proposed underpinning mechanism is likely to be modulated by a host of internal and external abiotic and biotic factors not addressed in our research so far. Both starch and sucrose levels in leaves are modulated by plant age and developmental stage, by the demand for substrates for respiration and biosynthesis within the leaf, and by progressive changes and short-term fluctuations in temperature, light intensity and quality, water and nutrient availability, CO_2_ concentration and symbiotic and pathogenic interactions with other organisms (e.g. Matheson and Wheatley, 1962; 1963; Chang, 1979; Egli et al., 1980; Hajibagheri and Flowers, 1985; Farrar and Williams, 1991; Schnyder, 1993; Brégard and Allard, 1999; Weise et al., 2006; Vriet et al., 2010; Thalmann et al., 2016; Thalmann and Santelia, 2017; Ruckle et al., 2017; Ning et al., 2018; Liang et al., 2019, Bruhn et al., 2022; Chu et al., 2022; Fan et al., 2026). It is thus unlikely that the underpinning mechanism alone will adequately describe carbohydrate dynamics in leaves in natural environments.

### Carbohydrate availability in grasses may be modulated by carbohydrate pools in leaf bases and sheaths

Although pooid grasses may resemble Arabidopsis and other eudicots with respect to the underpinning mechanism of control of diel carbohydrate turnover in mature leaves, they differ fundamentally in the manner in which carbohydrate is supplied from the leaf to other parts of the plant. Whereas in Arabidopsis and many other eudicots the fate of starch degraded at night in mature leaf blades appears relatively simple – it is converted to sucrose and used for maintenance and growth within the leaf or in sink tissues following export via the petiole – the same does not apply to mature leaf blades of pooid and many other grasses. Sucrose must pass from the mature region of the blade through its basal zone and then the leaf sheath prior to export to other plant organs. In grasses in general, these tissues are metabolically distinct from the fully expanded blade and from each other (e.g. Borland and Farrar, 1985; Schnyder et al., 1988; Solhaug, 1991; Williams et al., 1993; Guerrand et al., 1996; Shaikh et al., 2000; Pollock et al., 2003; Patton et al., 2007; Leegood, 2008; Verelst et al., 2013; Czedik-Eysenberg et al., 2016; Jensen and Wilkerson, 2017; Sadok et al., 2020). They have lower rates of photosynthesis than the blade, but usually contain substantial, discrete pools of starch, sucrose and often fructan that may display some diel turnover. In mature leaves, these pools may be largely derived from sucrose imported from the more distal parts of the blade. It seems likely that they act as buffers, damping down the very large diel fluctuation in the supply of sucrose from the distal regions of the blade leaf to permit a lower degree of diel fluctuation in carbohydrate supply to sink organs of the plant (Borland and Farrar, 1985; Shaihk et al., 2000; Czedik-Eysenberg et al., 2016). Feedback from these pools might also influence the nature and magnitude of diel fluctuations in carbohydrates in the distal part of the leaf blade.

In a preliminary experiment to examine the extent of diel carbohydrate turnover in leaf bases and sheaths in the conditions used for the time-course experiment, we measured starch and sucrose in blades, bases (basal 1 cm) and sheaths of the first, second and third most recently fully expanded leaf of 19-day-old wheat plants. In leaves in all three positions, starch and sucrose accumulated in leaf bases and sheaths during the day and was almost completely turned over during the night. These data suggest that leaf bases and sheaths as well as blades undergo diel carbohydrate turnover in wheat leaves and thus contribute to the night-time supply of carbohydrate to the rest of the plant (Figure S7).

In a very general sense, the role of the leaf base and sheath carbohydrate pools in grass species is analogous to the role of the arithmetic division mechanism in eudicot species with high starch turnover in leaves. Both mechanisms act to reduce the impact of diel and unexpected variation in daytime carbon assimilation on the availability of carbohydrate for growth and maintenance of the sink organs of the plant at night. The very different nature of the two mechanisms has profound implications for the impact of rhythmic and random environmental fluctuations on patterns of growth in grasses versus eudicots (e.g. Poiré et al., 2010; Matos et al., 2014; Müller et al., 2018) – a topic that is beyond the scope of this paper.

It is tempting to speculate that the large and robustly reproducible differences between pooid species in the ratio of sucrose to starch turnover in the mature regions of leaf blades are associated with differences in the size, nature and dynamics of the “carbohydrate buffer” in the blade base and the sheath. We found pronounced differences in the ratio of sucrose to starch turnover both within and between tribes and subtribes in the Pooideae. However - at least within our limited dataset - species in the same genus tended to have similar ratios. This might suggest that differences in the ratio of sucrose to starch turnover arose in the recent evolutionary past, in concert with a major proliferation and diversification of genera in the Pooideae believed to have occurred between nine and 15 million years ago (Zhang et al., 2022). It seems possible that diversification of leaf developmental patterns, morphology and relationships to sink organs that arose over this period was associated with diversification in the nature and capacity for carbohydrate storage and turnover in leaf bases and sheaths, affecting patterns of carbohydrate turnover in leaf blades. Further research is required to pinpoint the timing of divergence in carbohydrate turnover patterns, and to probe the selection pressures that drove it.

## MATERIALS and METHODS

### Plant material

Grass species were obtained as plants or seeds from in-house, seed bank and commercial sources, as shown in the table.

| Species | Source |
|---|---|
| *Calamagrostis brachytricha*<br><sup>2</sup>[*Calamagrostis arundinacea* (L.) Roth] | <sup>1</sup>Commercial |
| *Calamagrostis canescens* | <sup>3</sup>Royal Botanic Gardens, Millenium Seed Bank |
| *Calamagrostis scopulorum* | Royal Botanic Gardens, Millenium Seed Bank |
| *Agrostis capillaris* | Commercial |
| *Briza media* | <sup>4</sup>In house |
| *Anthoxanthum odoratum* | Commercial |
| *Koeleria macrantha* | Commercial |
| *Trisetum flavescens* | In house |
| *Lagurus ovatus* | Commercial |
| *Avena sativa* | In house |
| *Arrhenatherum elatius* | In house |
| *Echinaria capitata* | In house |
| *Holcus lanatus* | In house |
| *Phalaris coerulescens* | In house |
| *Festuca ovina* | Commercial |
| *Festuca pratensis*<br>[*Lolium pratense* (Huds.) Darbysh.] | Commercial |
| *Festuca rubra* | Commercial |
| *Lolium perenne* | In house |
| *Cynosurus cristatus* | In house |
| *Dactylis glomerata* | In house |
| *Poa trivialis* | In house |
| *Alopecurus geniculatus* | Commercial |
| *Alopecurus pratensis* | Commercial |
| *Deschampsia cespitosa* | Commercial |
| <sup>5</sup>*Triticum aestivum* | In house |
| *Secale cereale* | In house |
| *Secale iranicum* | In house |
| *Secale vavilovii* | In house |
| *Thinopyrum elongatum* | In house |
| *Thinopyrum intermedium* | In house |
| *Thinopyrum ponticum*<br>[*Thinopyrum obtusifolium* (DC.) Banfi] | In house |
| *Thinopyrum turcicum* | In house |
| *Hordeum jubatum* | In house |
| *Hordeum secalinum* | Commercial |
| <sup>6</sup>*Hordeum vulgare* | Commercial |
| *Brachypodium distachyon* | In house |
| *Brachypodium sylvaticum* | Commercial |
| *Stipa calamagrostis*<br>[*Achnatherum calamagrostis* (L.) PBeauv.] | Royal Botanic Gardens, Millenium Seed Bank |
| *Stipa conferta*<br>[*Timouria conferta* (Poir.) Sennikov] | Royal Botanic Gardens, Millenium Seed Bank |
| *Melica altissima* | Commercial |
| *Lygeum spartum* | Royal Botanic Gardens, Millenium Seed Bank |

<sup>1</sup>Commercial sources were: The Plantsman’s Preference (www.plantpref.co.uk); Jelitto Seeds (www.jelitto.com); Plant World Seeds (www.plant-world-seeds.com); Chiltern Seeds (www.chilternseeds.co.uk); Emorsgate Seeds (www.wildseed.co.uk).
<sup>2</sup>Species names in this paper are those under which the material was acquired. In five cases, these are not the currently accepted names on the definitive Plants of the World Online database (POWO; Plants of the World Online \| Kew Science). The POWO accepted names are given in square brackets. None of the name differences affect the phylogenetic relationships of the species. Where the POWO generic name is different from the name used in this paper, the two names are homotypic synonyms.
<sup>3</sup>www.kew.org/science/collections-and-resources/research-facilities/millennium-seed-bank
<sup>4</sup>In-house resources were the John Innes Centre Germplasm Resource Unit (www.jic.ac.uk/research-impact/germplasm-resource-unit/), the collection of Dr. Matthew Moscou (The Sainsbury Laboratory, Norwich), and collections of other John Innes Centre staff.
<sup>5</sup>cv Cadenza
<sup>6</sup>cv Tipple

### Plant growth conditions

For glasshouse experiments, plants were grown in individual pots in soil-based compost, either from seed or from divisions of stock plants where seed set and/or viability were problematic. Plants had 10 h natural daylight (controlled by automated movement of shelves in and out of a dark shed) and ambient temperature conditions. For controlled environment experiments, all plants were grown from seed in individual pots in soil-based compost. Plants were grown at 20°C, with the day lengths and light intensities (LED lights) specified in the text and figures. For metabolite analyses, the central 1-cm section of the youngest fully expanded blade was taken from young, non-flowering, non-tillering plants. Depending on the growth rate of the species, plants were between about two and seven weeks old at the time of harvest.

### Metabolite analyses

Leaf blade sections were harvested rapidly onto dry ice and stored at -70°C until analysis. Homogenisation, perchloric acid extraction, extract preparation and enzymatic assays of starch and sucrose were as described by Martins et al. (2013).

Enzymatic assays to test for the presence of fructans were performed on the soluble extracts used for sucrose assays. For assays with sucrase, lyophilised sucrase (Megazyme: www.megazyme.com) was dissolved in 100 mM maleic acid buffer (pH 6.5) and 1.4 units in 100 μl was incubated with 100 μl of a two-fold dilution of soluble extract overnight. Controls contained maleic acid buffer without enzyme. Samples and controls were then assayed enzymatically for glucose and fructose. For assays with fructanase, 10 units of fructanase (Megazyme, liquid) in 100 μl 100 mM Na acetate buffer (pH 4.5) was incubated with a two-fold dilution of soluble extract for 2 h at 37°C. Controls contained Na acetate buffer without enzyme. Samples and controls were then assayed enzymatically for glucose and fructose. Further details of checks and controls for these assays are given in Figures S2 and S3.

High-pressure anion exchange chromatography (HPAEC) measurements of fructans were made with a Carbopac PA1 column on a Thermo Fisher Scientific (www.thermofisher.com) Dionex ICS-5000 system with a DC detector and Chromeleon software. Samples of 0.5 ml of soluble extract were diluted two-fold and passed through sequential 1.5-ml columns of Dowex 1 and Dowex 50 ion exchange resins, followed by 1 ml water. Eluates were freeze-dried then suspended in appropriate volumes of buffer A (100 mM NaOH). Samples of 5 μl were injected and eluted at a flow rate of 0.25 ml min^-1^ with a gradient from 100% buffer A to 40% buffer A, 60% buffer B (100 mM NaOH, 600 mM Na acetate) over 31 min.

## ACKNOWLEDGEMENTS

This work was funded by postdoctoral grants from the Swiss National Science Foundation to MT, and by Institute Strategy Funding from the John Innes Centre to AMS. We are grateful to Matthew Moscou (formerly at The Sainsbury Laboratory, Norwich, UK), the Millennium Seed Bank (Royal Botanic Gardens, Wakehurst Place, UK), Noam Chayut (John Innes Centre Germplasm Resource Unit) and other JIC colleagues for making grass seeds available for this project. We thank Martin Rejzek (John Innes Centre Chemistry Platform), Marilyn Pike, Brendan Fahy and staff of John Innes Centre Horticultural Services for support and advice. We benefitted from many discussions with David Seung and lab members (John Innes Centre) and John Lunn (Max-Planck Institute for Molecular Plant Physiology, Potsdam). Mark Stitt (Emeritus Director, Max-Planck Institute for Molecular Plant Physiology, Potsdam) and David Seung gave invaluable feedback on the manuscript.

## CONTRIBUTIONS

MT and AMS designed the project, and MT carried it out. AMS wrote the manuscript, with input from MT.

## Supplementary data

**Figure S1.** Percentage of end-of-day starch/sucrose remaining at the end of the night.

**Figure S2.** Checks for fructans in extracts of leaves grown in 10 h light, 10 h dark in a glasshouse.

**Figure S3.** Checks for fructans in extracts of leaves grown in high light and long days.

**Figure S4.** Total amount of carbohydrate turned over during the night in leaves of six grass species.

**Figure S5.** Examples of HPAEC analyses for fructan contents.

**Figure S6.** Analysis of the decline in sucrose content in leaves at night.

**Figure S7.** End-of-day and end-of-night sucrose and starch contents of wheat leaf blades, bases and sheaths.

## Notes

### Competing Interest Statement

The authors have declared no competing interest.

### Summary of Updates

This version of the manuscript has been revised to add a small number of recent references and adjust the text accordingly. There are also minor changes to text and figures intended improve clarity and readability. These have no material effect on the results and conclusions presented in the original version of the manuscript.

