## Supplementary data for "Night-time control of carbohydrate availability in grasses differs radically from that in Arabidopsis"

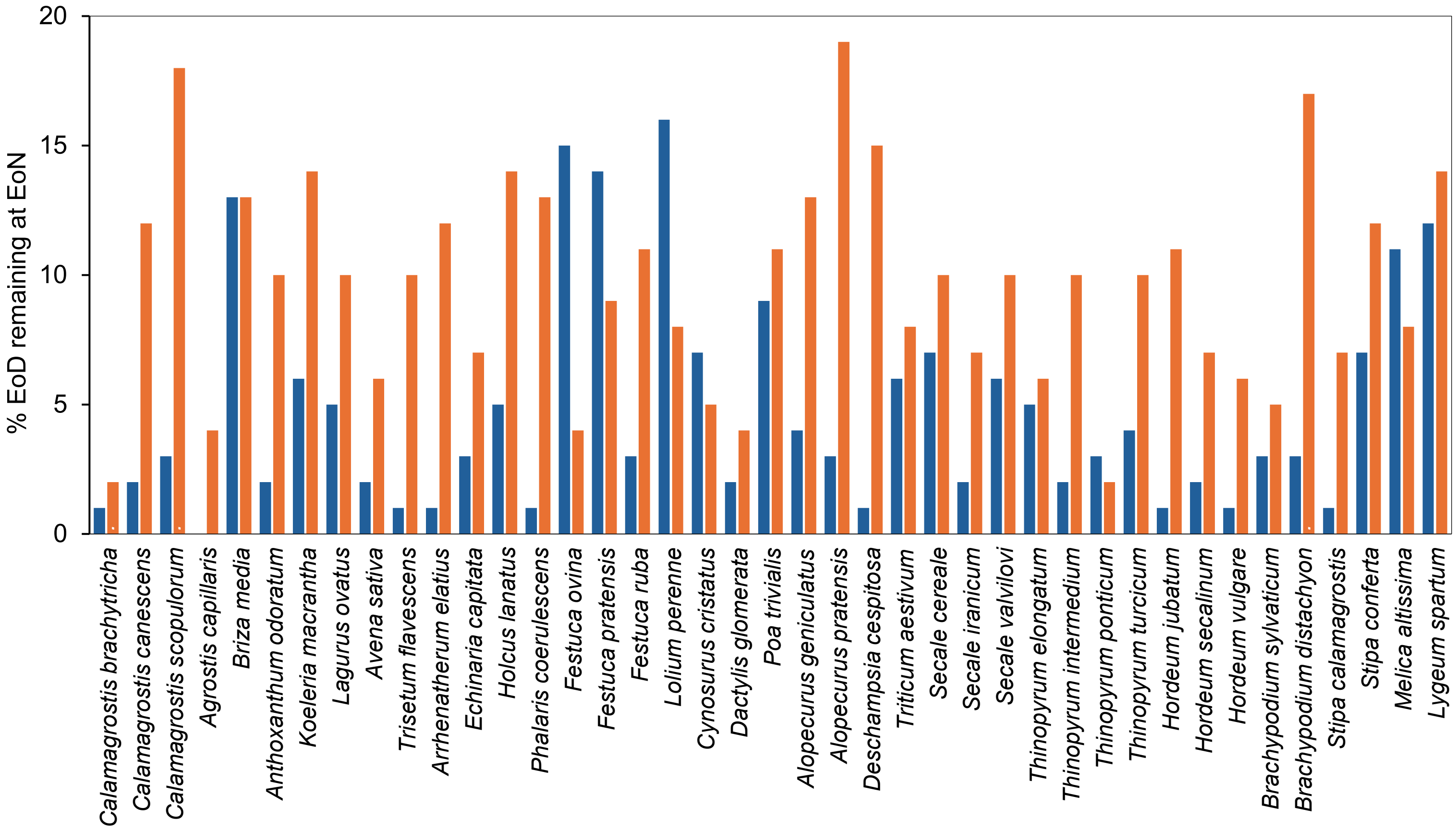


**Figure S1.**

**Percentage of end-of-day starch/sucrose remaining at the end of the night**.

Values were calculated from data in Figure 1. Orange bars: sucrose. Blue bars: starch.

For each species, values are: $\frac{\text{EoD-EoN}}{\text{EoD}} \times 100$

.

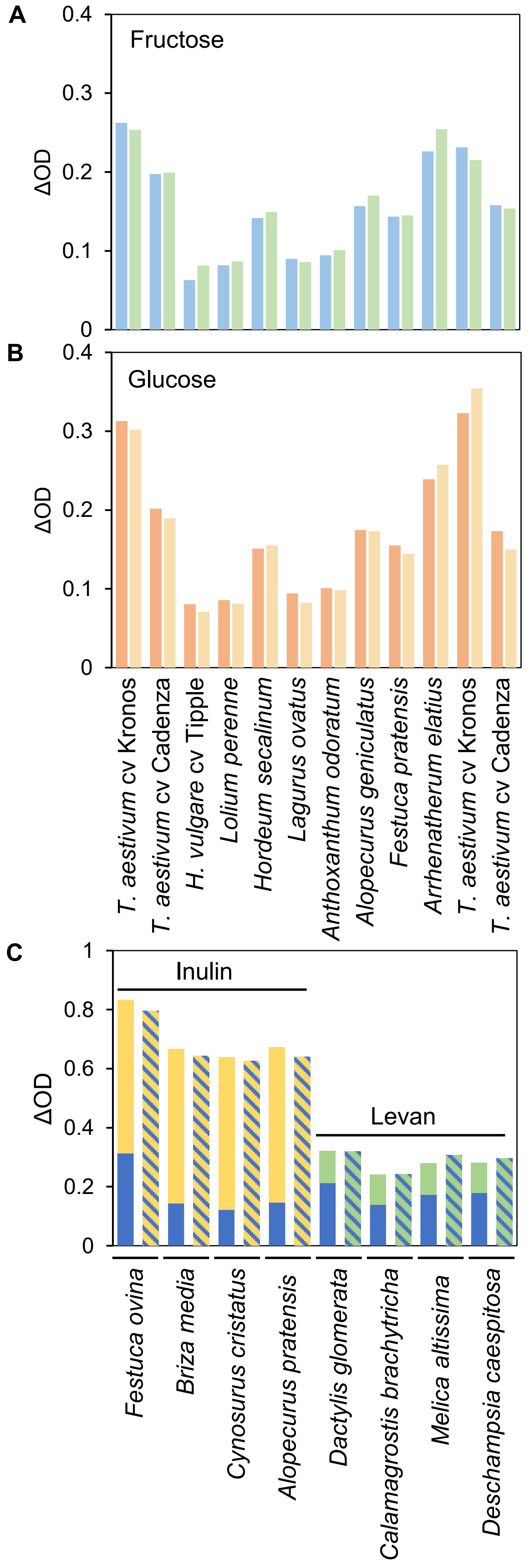


**Figure S2.**
**Checks for fructans in extracts of leaves grown in 10 h light, 10 h dark in a glasshouse**. Youngest fully expanded leaves were harvested at the end of the light period. Bars represent means of 3 technical replicates on individual extracts randomly chosen from those used for the measurements presented in Figure 2. **A** and **B** compare the hexose products of invertase digestion with those of sucrase and fructanase digestions. **C** shows a recovery experiment to check the impact of plant extracts on fructanase digestions of inulin and levan standards. **A** Fructose produced by digestion with sucrase (blue bars) or invertase (green bars). **B** Glucose produced by digestion with fructanase (orange bars) or invertase (pale orange bars). Two independent extracts were sampled for *Triticum aestivum* cv Kronos and cv Cadenza. Assays on extracts from 8 further species also showed no difference between hexose yields from invertase and fructanase digestions. These species were: *Festuca ovina*, *Briza media*, *Cynosurus cristatus*, *Alopecurus pratensis*, *Dactylis glomerata*, *Calamagrostis brachytricha*, *Melica altissima*, *Deschampsia caespitosa*. **C** For each species, the left bar shows the amount of hexose produced in a fructanase digestion of a plant extract (lower, blue section of bar), and the amount of hexose produced in a fructanase digestion of a fructan standard (upper yellow (inulin) or green (levan) section of bar). The right bar for each species (yellow or green with blue stripes) shows the amount of hexose produced in a fructanase digestion of a mixture of the extract and fructan standard for which the outcomes of separate digestions are shown in the left bar.

Taken together, these results indicate that extracts of leaves grown in 10 h light, 14 h dark do not contain small fructo-oligosaccharides or larger fructans.

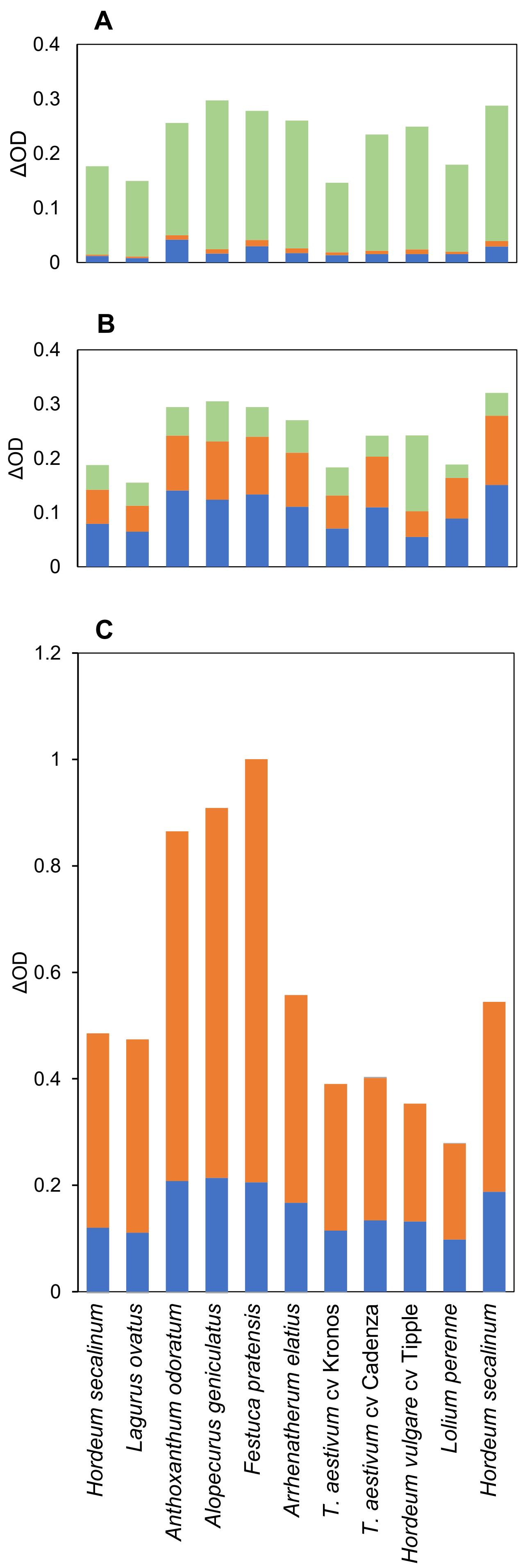


**Figure S3.**
**Checks for fructans in leaves grown in high light and long days.** Plants were grown in a CER with 18 h light (400 µmol PAR $m^{-2}$ $s^{-1}$), 6 h dark, at 20°C. Youngest fully expanded leaves were harvested at the end of the light period. Values are means of 3 technical replicates on individual extracts randomly chosen from those used in Figure 4 or on extracts made from plants grown at the same time in the same conditions. **A**, **B** and **C** compare the hexose products of invertase digestion (**A**) with those of sucrase+invertase (**B**) and fructanase+invertase (**C**) digestions. **A** The lower two sections of the bars show amounts of free glucose (blue) and fructose (orange) in extracts prior to any digestion. The upper, green sections show the amount of hexose (glucose+fructose) produced by invertase digestion. **B** The lower two sections of the bars show total amounts of glucose (blue) and fructose (orange) present in the extracts shown in **A**, after sucrase digestion. The upper green sections show the additional hexoses (glucose+fructose) released by invertase digestion following sucrase digestion. **C** The blue and orange sections of the bars show the amounts of glucose (blue) and fructose (orange) present in the extracts shown in A and B, after fructanase digestion. Invertase digestion of extracts digested with fructanase did not release any additional hexose.

These results indicate that only small amounts of free hexoses are present in leaves at EoD (**A**). The fact that sucrase+invertase digestion released more hexose than sucrase digestion alone indicates that small fructo-oligosaccharides, susceptible to digestion by invertase but not sucrase, are present in the extracts (**B**). The production by fructanase digestion of much more hexose, largely fructose, than produced by sucrase+invertase digestion (**C**) indicates that the extracts contain large fructans not susceptible to invertase digestion as well as small fructo-oligosaccharides.

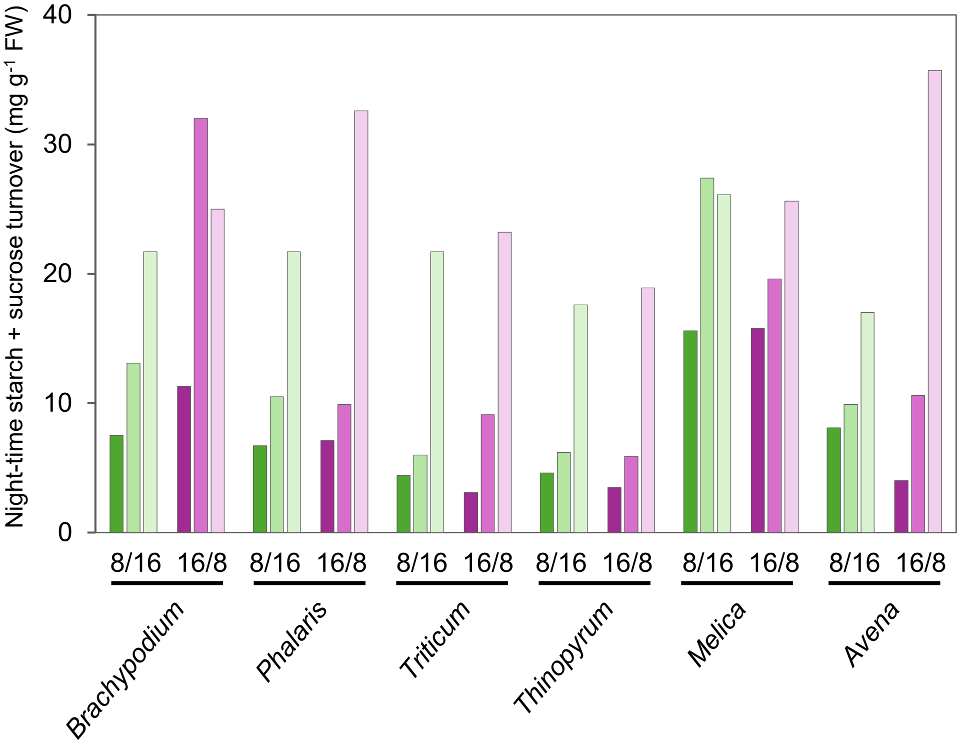


**Figure S4.**

**Total amount of carbohydrate turned over during the night in leaves of six grass species.**

Total carbohydrate is the sum of sucrose and starch contents. Data were taken from Figure 5B, and are (EoD sucrose+starch) – (EoN sucrose+starch) for plants grown in short and long days (8/16 and 16/8 h, respectively), with 100, 200 or 400 µmol quanta PAR $m^{-2}$ $s^{-1}$ (dark, mid and light colours respectively, green for short days, purple for long days).

**Figure S5.**
**Examples of HPAEC analyses for fructan contents.**
**A** Injection of standard solutions of sucrose and the fructo-oligosaccharides nystose and kestose showed that detector response was linear over a 10-fold concentration range. Points (mostly overlapping) represent two or three technical replicates at each concentration. **B** Elution profiles of standard solutions showed clean separation of hexoses, sucrose and fructo-oligosaccharides. **C**. Examples of profiles obtained from desalted extracts of *Triticum* leaves. Top: Plants were grown in high light and long days and harvested at the end of the light period. Middle: As for Top but harvested at the end of the night. Bottom: Plants were grown in low light with 12/12 days and harvested at the end of the day. The complex profiles putatively representing fructo-oligosaccharides are underlined in red.

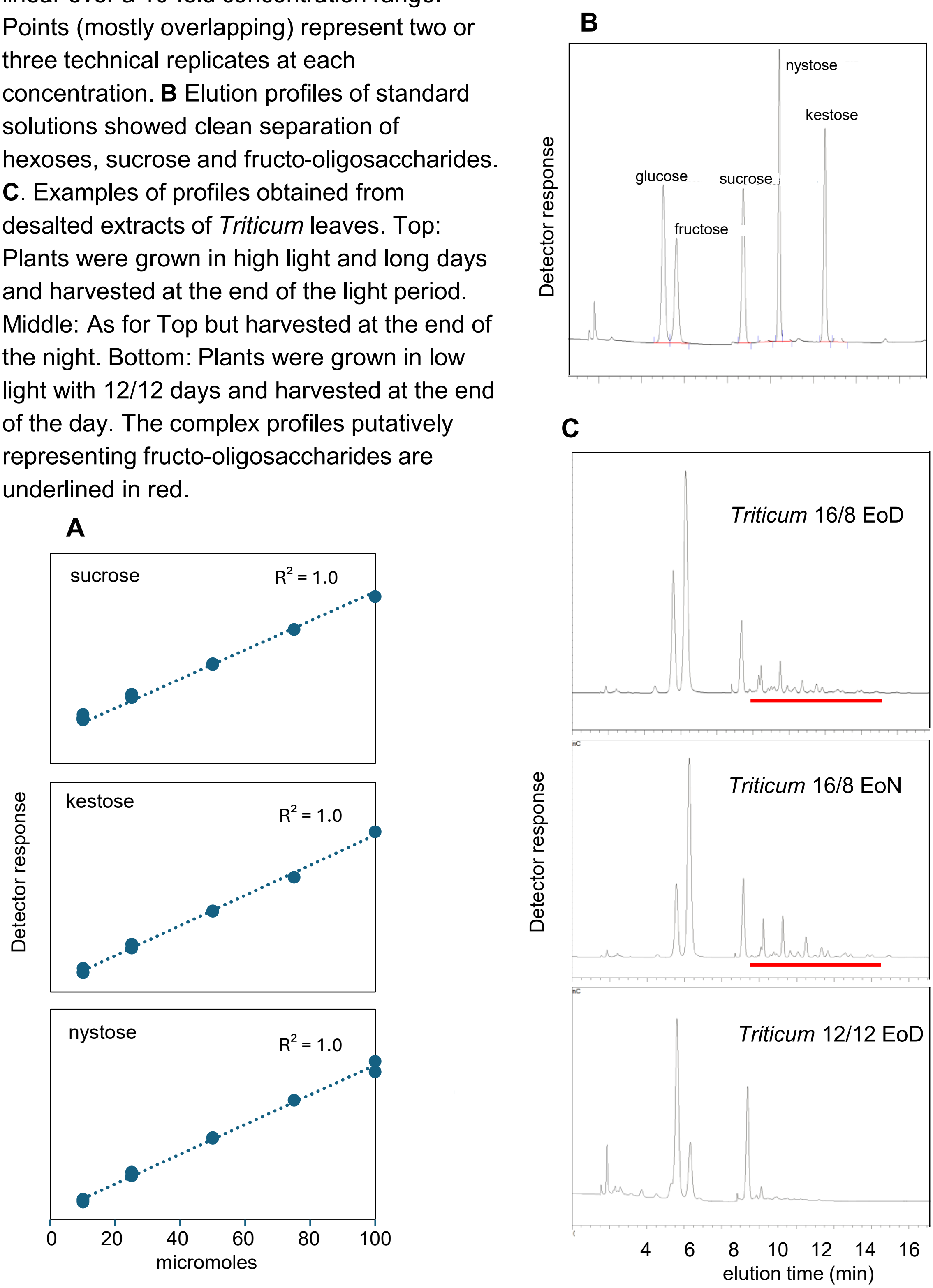

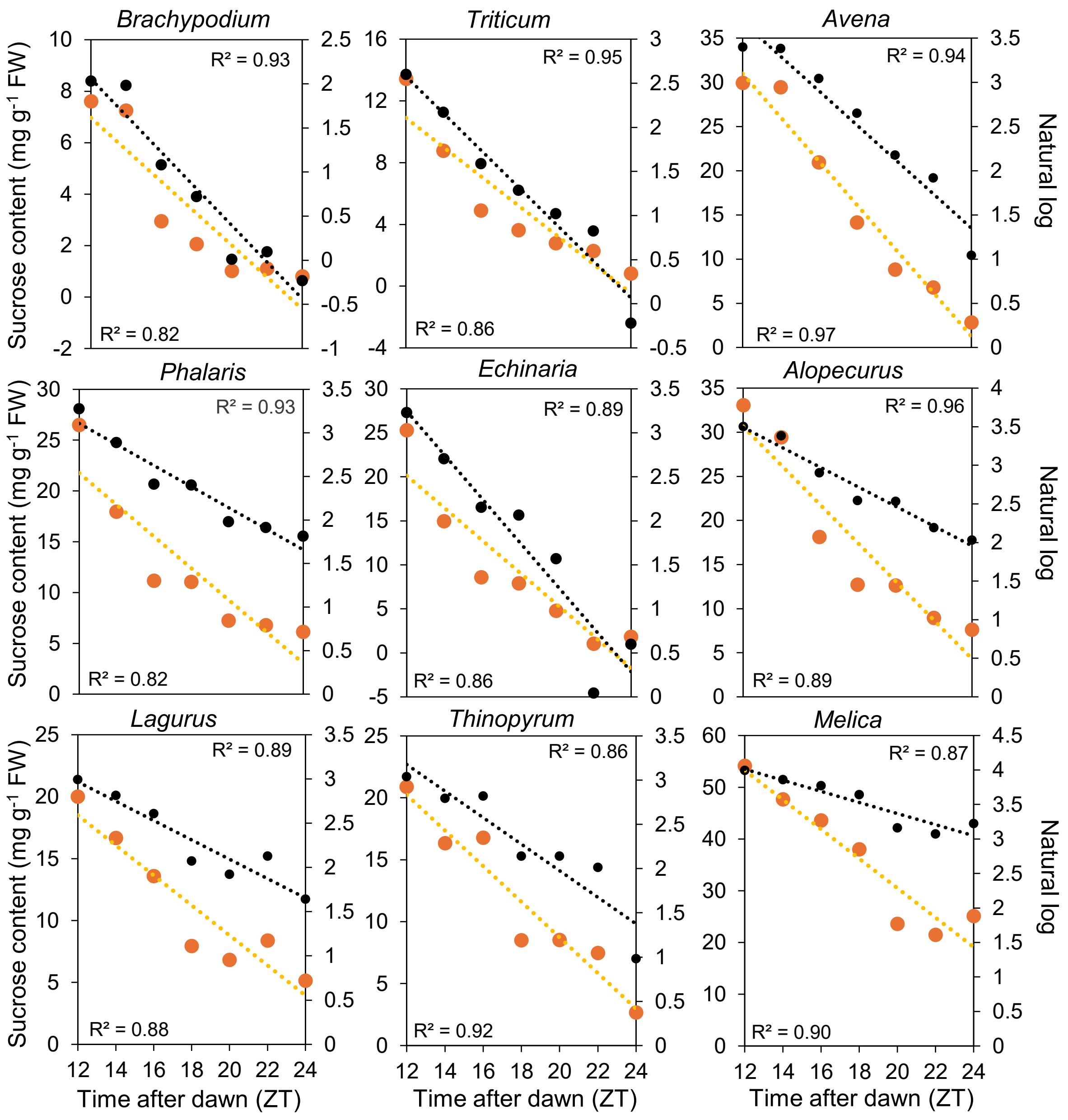


**Figure S6.**

**Analysis of the decline in sucrose content in leaves at night.**

Data from Figure 7 for sucrose content over the night period (ZT12 to ZT24; orange symbols) are plotted with the best fit to a straight line (orange dashed line) and the best fit of natural logs of these values to a straight line (right axis; black dashed line is line of best fit). $R^2$ values are given at the bottom left (orange line) and top right (black line) of the graph.

The graphs show that, in general, the data do not allow clear distinctions between linear and exponential declines in sucrose contents.

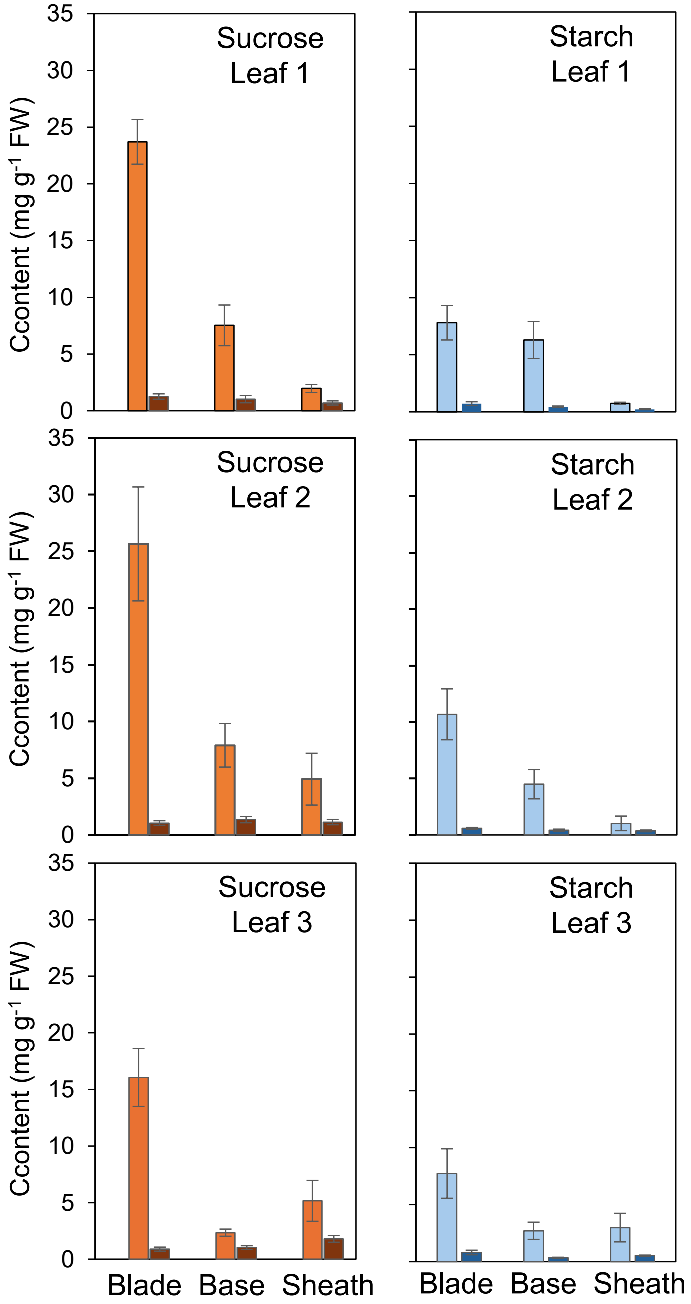


**Figure S7.**

**End-of-day and end-of-night sucrose and starch contents of wheat leaf blades, bases and sheaths**.

The central 1-cm section of the blade, the basal 1 cm of the blade, and the sheath of the first, second and third most recently expanded leaf (Leaf 1, Leaf 2, Leaf 3) of 19-day-old plants grown in 12 h light, 12 h dark and 400 µmol quanta PAR $m^{-2}$ $s^{-1}$ in a CER were sampled at end of the day or the end of the night. Values are means ±SE of measurements on samples from 6 different plants. Panels show sucrose and starch at the end of the day (orange and light blue bars respectively) and the end of the night (brown and dark blue bars respectively).
